# Transcriptional control and exploitation of an immune-responsive family of plant retrotransposons

**DOI:** 10.1101/206813

**Authors:** Jérôme Zervudacki, Agnès Yu, Delase Amesefe, Jingyu Wang, Jan Drouaud, Lionel Navarro, Angélique Deleris

## Abstract

Mobilization of transposable elements (TEs) in plants has been recognized as a driving force of evolution and adaptation, in particular by providing genes with regulatory modules that impact their transcription. In this study, we employed an *ATCOPIA93* Long terminal repeats (LTR) promoter-*GUS* fusion to show that this retrotransposon behaves like an immune-responsive gene during plant defense in Arabidopsis. We also showed that the reactivation of the endogenous *ATCOPIA93* copy *“EVD*”, in the presence of bacterial stress, is not only negatively regulated by DNA methylation but also by Polycomb-mediated silencing—a mode of repression typically found at protein-coding and microRNA genes. Interestingly, one of the *ATCOPIA93*-derived soloLTRs is located upstream of the disease resistance gene *RPP4* and is devoid of either DNA methylation or H3K27m3 marks. Through loss-of-function experiments, we demonstrated that this soloLTR is required for proper expression of *RPP4* during plant defense, thus linking the responsiveness of *ATCOPIA93* to biotic stress and the co-option of its LTR for plant immunity.

## Introduction

TEs are repeated sequences that can potentially move and multiply in the genome. Their mobilization has been recognized as a driving force of evolution and adaptation in various organisms, in particular by providing genes with regulatory modules that can create or impact transcriptional programs (Chuong *et al*, 2016). The study of TE regulation is thus important in order to understand both the conditions for their transposition but also their influence, as full-length or truncated elements, on nearby gene regulation. This role in *cis* has been demonstrated in plants by artificially inducing insertions that confer gene regulation, *e.g.*, the rice TE *mPing* (Naito *et al*, 2009) or the Arabidopsis TE *ONSEN* (Ito *et al*, 2011). However, the causal link between *cis*-regulatory properties of TEs and established expression patterns of nearby genes, requires loss-of-function experiments and has rarely been demonstrated (Chuong *et al*, 2016).

TE *cis*-regulatory effects can either be genetic in nature, such as when the TE contains regulatory motifs, or epigenetic through recruitment of dimethylation of histone 3 lysine 9 (H3K9m2) and cytosine DNA methylation, which are hallmarks of transposon control. DNA methylation in Arabidopsis is carried out by four pathways. While METHYLTRANSFERASE1 (MET1) maintains CG methylation (Kankel *et al*, 2003)(Cokus *et al*, 2008; Lister *et al*, 2008), CHROMOMETHYLASE2 and 3 (CMT2 and CMT3) maintain CHG methylation (where H is any base pair but not a G) (Zemach *et al*, 2013; Stroud *et al*, 2014). The maintenance of CHH methylation requires either CMT2 for long heterochromatic repeats or RNA-directed DNA methylation (RdDM) mediated by DOMAINS REARRANGED METHYLASE 2 (DRM2) and accompanying small RNA machinery (Cao & Jacobsen, 2002; Chan, 2004; Stroud *et al*, 2013; Zemach *et al*, 2013). In addition, the SNF2 family chromatin remodeler DECREASED DNA METHYLATION 1 (DDM1) is necessary for heterochromatic DNA methylation in all cytosine sequence contexts (Jeddeloh *et al*, 1999; Zemach *et al*, 2013; Stroud *et al*, 2013). Furthermore, DNA methylation and H3K9m2 are mechanistically interconnected and, as a result, are largely co-localized throughout the genome (Du *et al*, 2015). Importantly, this histone and cytosine marking can also impact the nearby genes which then become epigenetically controlled because of the inhibitory effect of DNA and H3K9 methylation on promoter activity (Lippman *et al*, 2004; Liu *et al*, 2004; Huettel *et al*, 2006; Gehring *et al*, 2009). In addition, in both plants (Mathieu *et al*, 2005; Deleris *et al*, 2012; Weinhofer *et al*, 2010) and animals (Reddington *et al*, 2013; Saksouk *et al*, 2014; Basenko *et al*, 2015; Walter *et al*, 2016), the removal of DNA methylation at some

TEs leads to H3K27 trimethylation (H3K27m3), an epigenetic mark deposited and interpreted by Polycomb Group (PcG) proteins, which normally target and silence protein coding genes that are often developmentally important (Förderer *et al*, 2016). Thus, there exists a potential for this alternative repression system to mediate silencing of TEs, but it has not been fully explored in plants, except in the endosperm, a nutritive and terminal seed tissue that is naturally DNA hypomethylated (Weinhofer *et al*, 2010; Moreno-Romero *et al*, 2016).

In accordance with the epigenetic control mediated by DNA and histone H3K9 methylation, mutations in DNA methylation pathway genes lead to reactivation of various subsets of TEs; however, these defects in chromatin regulation are not always sufficient for TE expression, and the activation of specific signaling pathways is sometimes needed. This has been exemplified by the Arabidopsis Long-terminal repeats (LTR)-retrotransposon *ONSEN*, which was shown to be reactivated after heat-stress, in wild-type plants and independently from a loss of DNA methylation in this context (Ito *et al*, 2011; Cavrak *et al*, 2014). In addition, *ONSEN* was not expressed in unstressed RdDM-defective mutants (nor in *ddm1*), but its induction was enhanced in RdDM-defective mutants subjected to heat stress (Ito *et al*, 2011). To our knowledge, *ONSEN* is the only described example of a TE which expression is modulated by DNA methylation during stress response. Thus, it is important to characterize other TEs that exploit plant signaling, in order to gain a deeper understanding of the connection between biotic/abiotic stresses and transposon activation, and how epigenetic silencing pathways exert their influence on this relationship.

In this study, we unravel the responsiveness of another family of Arabidopsis retroelements, *ATCOPIA93* (Mirouze *et al*, 2009; Marí-Ordóñez *et al*, 2013), during PAMP-triggered immunity (PTI). PTI is defined as the first layer of active defense against pathogens and relies on the perception of evolutionary conserved Microbe- or Pathogen-Associated Molecular Patterns (MAMPs or PAMPs) by surface receptors (Boutrot & Zipfel, 2017). *ATCOPIA93* is a low-copy, evolutionary young family of LTR-retroelements, which is tightly controlled by DNA methylation, in particular CG methylation (Mirouze *et al*, 2009a). The family representative *EVD* (*AT5G17125*) was found to transpose in *ddm1* after eight generations of inbreeding (Tsukahara *et al*, 2009) as well as in genetically wild-type epigenetic recombinant lines (epiRILs) derived from crosses between wild-type and *met1* (Mirouze *et al*, 2009, Marí-Ordóñez *et al*, 2013) or *ddm1* (Marí-Ordóñez *et al*, 2013). *EVD* is 99.5% identical in sequence to the pericentromeric *ATR* (*AT1G34967*), which is predicted to encode a polyprotein but does not seem to be active in the latter conditions (Mirouze *et al*, 2009). Here, we first took advantage of an unmethylated *ATCOPIA93* LTR-*GUS* fusion that we used as a reporter of promoter activity, since LTRs of retroelements contain *cis*-regulatory sequences that can recruit RNA Pol II (Chuong *et al*, 2016). We showed that this LTR exhibits the hallmarks of a promoter of an immune-responsive gene in the absence of epigenetic control. Accordingly, the corresponding methylated endogenous *ATCOPIA93* retroelements, *EVD* and *ATR,* were significantly more reactivated after PAMP-elicitation in a DNA hypomethylated background, in *met1* or *ddm1* mutants, than in the wild type. Interestingly, we demonstrated for the first time in wild-type plant vegetative tissues, a second layer of control of TE expression mediated by Polycomb silencing. Importantly, we showed that H3K27m3 co-exists with DNA methylation at *EVD* sequences but not at *ATR*, leading to a differential negative control between these two copies during immunity. Furthermore, we were able to test the implications of these findings for the regulation of the immune response. We identified an *ATCOPIA93*-derived soloLTR, unmethylated and not marked by H3K27m3, upstream of the *RECOGNITION OF PERONOSPORA PARASITICA 4* (*RPP4)* disease resistance gene. By inducing the genetic loss of this soloLTR, we could show that it has been co-opted for the proper expression of *RPP4* during basal defense triggered by unrelated PAMPs and plays a role as regulatory “enhancer” element. Thus, we established a link between the responsiveness of a TE to biotic stresses and the co-option of its derived soloLTR for plant immunity, where the repressive epigenetic modifications controlling the full-length active elements are absent on the derived regulatory sequence.

## Results

### *ATCOPIA93*-LTR::*GUS* transcriptional fusion behaves as a canonical immune-responsive gene

An *ATCOPIA93*-LTR::*GUS* construct —comprising the full *EVD/ATR* LTR upstream of a sequence encoding a GUS protein (schema Fig 1A, EV1A) — was transformed into the Arabidopsis wild type reference accession Columbia (Col-0), initially to serve as a reporter of DNA methylation levels as previously reported for a Gypsy-type retroelement LTR (Yu *et al*., 2013). Unexpectedly, the LTR::*GUS* transgenes were not methylated in any of the transgenic lines obtained (Fig 1A, Fig EV1B-C). Thus, instead, we used the LTR::*GUS* as a reporter of *ATCOPIA93* promoter activity that we could exploit to assess *ATCOPIA93* responsiveness during PTI in the absence of DNA methylation-mediated control. Plants containing the LTR::*GUS* transcriptional fusion were further elicited with either flg22 (a synthetic peptide corresponding to the conserved N-terminal region of bacterial flagellin that is often used as a PAMP surrogate) (Zipfel *et al*, 2004; Boutrot & Zipfel, 2017), or *Pto*Δ28E (a non-pathogenic *Pseudomonas syringae* pv. *tomato* DC3000 (*Pto* DC3000) in which 28 out of 36 effectors are deleted (Cunnac *et al*, 2011), and the accumulation of *GUS* mRNA and protein was monitored over a 24 hour time course. In water-treated plants, at 24 hours post-infiltration (hpi), there was barely any GUS staining, indicating that the activity of the *ATCOPIA93* LTR promoter is weak in this condition. By contrast, in both flg22 and *Pto*Δ28E treatments, an intense GUS staining was observed at 24 hpi (Fig 1B, top panel, Fig EV1D). This was associated with progressive GUS protein accumulation over the time-course, until it reaches a plateau (Fig 1B, bottom panel). At all the time points analyzed, the *GUS* expression was generally stronger in response to *Pto*Δ28E than flg22 and thus we focused on the *Pto*Δ28E bacterial elicitor for the rest of the study. By analyzing GUS mRNA levels, we could then show that the *GUS* induction was transient, similarly as an immune-responsive gene induced rapidly during PTI such as *WRKY29* (Asai et al., 2002) (Fig 1C, Fig EV1E). In addition, treatment with the virulent wild-type *Pto* DC3000 strain, which can inject type III effectors into the host cell, resulted in a partially compromised induction of GUS protein accumulation compared to a similar inoculum of *Pto*Δ28E (Fig 1D), suggesting that some bacterial effectors suppress the *LTR* responsiveness during PTI. This transcriptional behavior is reminiscent of typical PTI-induced genes whose induction is impaired by bacterial effectors that have evolved to suppress different steps of PTI to enable disease (Asai & Shirasu, 2015). Together, these data show that the *ATCOPIA93* LTR::*GUS* transgene behaves like a canonical immune-responsive gene, which is transcriptionally reactivated during PTI and whose induction is suppressed by bacterial effectors. In support of this, we noticed in the LTR sequence the presence of two putative W-box elements, i.e. DNA sequences with the C/TTGACC/T (A/GGTCAA/G) motif, which are the cognate binding sites for WRKY transcription factors that are known to orchestrate transcriptional reprogramming during PTI (Rushton *et al*, 2010) (Fig EV1A). Importantly, we found that these two putative W-box elements are functional as the *Pto*Δ28E-mediated transcriptional induction of *GUS* was lost in part or entirely when W-box 1 and W-box 2 were mutated, respectively (Fig 1E).

**Figure 1.**
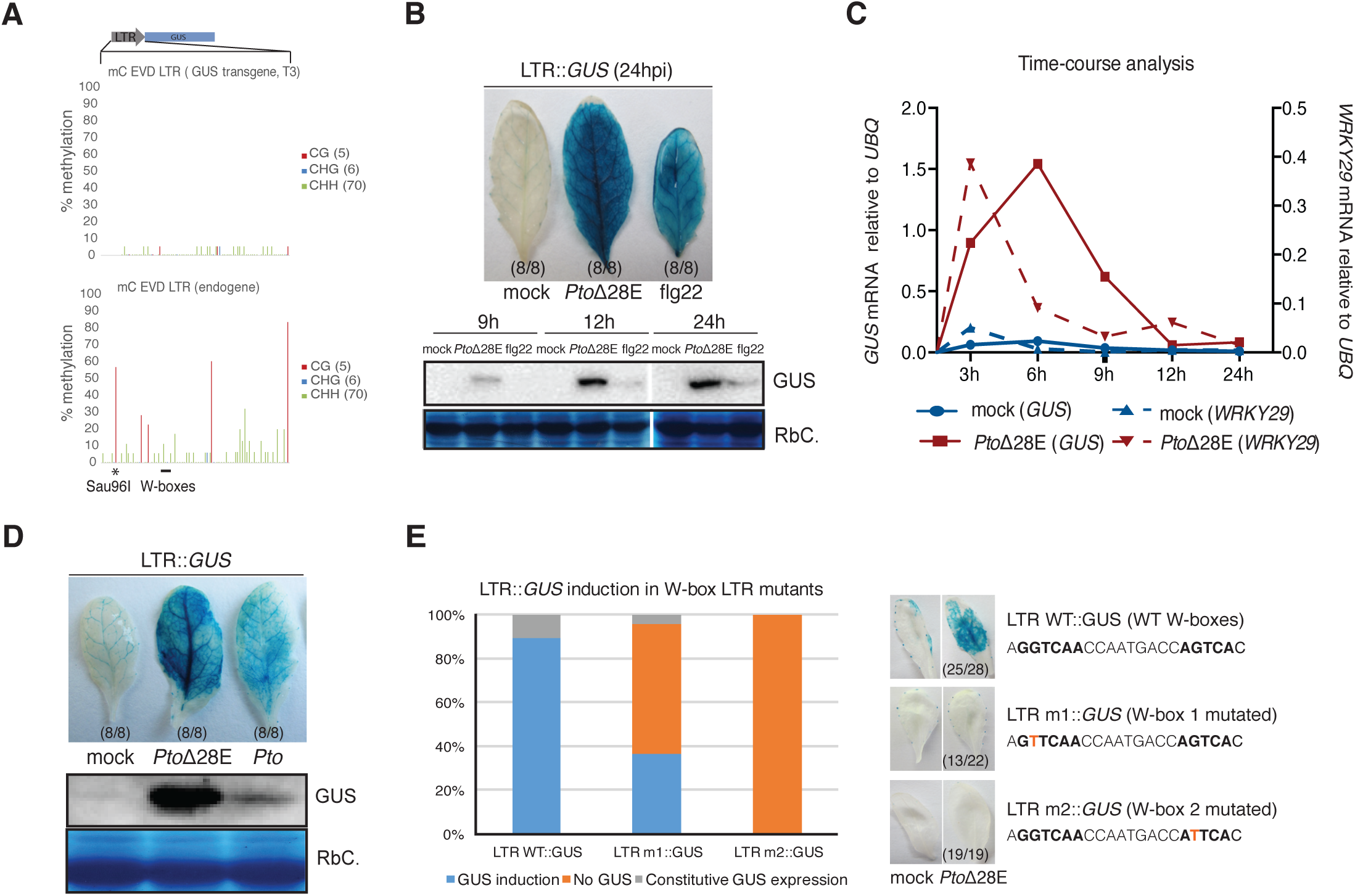
*ATCOPIA93* LTR::*GUS* transcriptional fusion behaves like a canonical immune responsive gene. **A.** Cytosine methylation analyzed by bisulfite-sequencing at the LTR::*GUS* transgene. Genomic DNA of a pool of LTR::*GUS* transgenic plants (one T3 line: T3#12, four plants, two rosette leaves each) was treated with sodium bisulfite, amplified with primers specific for the LTR contained in the LTR::*GUS* construct and cloned for sequencing (19 clones). The endogenous LTR of *ATCOPIA93* EVD was sequenced as a positive control (17 clones). The analysis of another T3 line led to the same results (Fig EV1B). The percentage of methylated cytosines is indicated by vertical bars. The number of CG, CHG and CHH sites is indicated on the right. This result was also reproduced in various T3 and T1 lines by a Sau96I methylation-sensitive assay analyzing the first CG site (black asterisk) (Fig EV1C). **B.** Accumulation of GUS protein detected in response to bacterial elicitors of basal immunity. Upper panel: representative pictures of leaves infiltrated with water (mock), *Pto* DC3000 deleted of 28 effectors (*Pto*Δ28E) at 2.10^8^ colony-forming unit per ml (cfu/ml) or 1 μM of flg22, and incubated with GUS substrate 24 hours post-infiltration (24hpi). The number of leaves showing this representative phenotype is indicated into brackets. The T3 line LTR::*GUS* #12 (used for the remainder of the study) is shown here but two additional homozygous T3 lines displayed the same phenotype and are presented in Fig EV1D. Lower panel: Western blot analysis over a 24h time-course; RbC: Rubisco. Three to four plants (two leaves per plant) were infiltrated for each condition and time point, and leaves pooled by condition and time-point before extracting the proteins. Samples derived from the same experiment, and gels and blots were processed in parallel. This experiment was repeated twice with similar results. **C.** Time-course analysis of *GUS* mRNA (plain lines) and PTI-marker *WRKY29* mRNA (dashed lines) by RT-qPCR. Leaves were infiltrated with water (mock), or *Pto*Δ28E bacteria at 2.10^8^ cfu/ml; two similar leaves of three to four plants were pooled by condition and by time-point after infiltration (as in B) before extracting the RNA subjected to RT-qPCR. Values are relative to the expression of the *UBIQUITIN* gene (*At2g36060*). This experiment was repeated twice independently and another independent experiment is shown in Fig EV1E. **D.** Accumulation of GUS protein detected in response to virulent *Pto* DC3000 versus *Pto*Δ28E. Upper panel: representative pictures of leaves infiltrated with water (mock), effectorless (*Pto*Δ28E) and virulent (*Pto*) bacteria *Pto* DC3000, both at 1.10^7^ cfu/ml, and incubated with GUS substrate at 24hpi. The number of leaves showing this representative phenotype is shown into brackets. Lower panel: Western blot analysis of the GUS protein accumulated at 9hpi; RbC: Rubisco. Two similar leaves of three to four plants were pooled by condition and time-point before extracting the proteins. This experiment was repeated twice with similar results. **E.** Activation of the GUS expression upon *Pto*Δ28E elicitation in LTR::*GUS* plants with mutated W-boxes. Experiments were performed on 28, 22, and 19 primary transformants for the LTR::*GUS* WT, m1 and m2 constructs respectively; the point-mutations introduced are depicted on the right (W-boxes are the sequences in bold). Mock and *Pto*Δ28E (at 2.10^8^ cfu/ml) treatments were performed on four similar leaves of each individual transformant and GUS staining performed 24 hours later on two leaves. One representative picture (into brackets is the number of plants showing this phenotype) for one primary transformant is shown for each construct with each treatment. Plants were classified into three categories: normal GUS induction, loss of GUS induction, constitutive GUS expression and percentages of plants belonging to each category over the total number of plants tested were calculated.

**Figure EV1.**
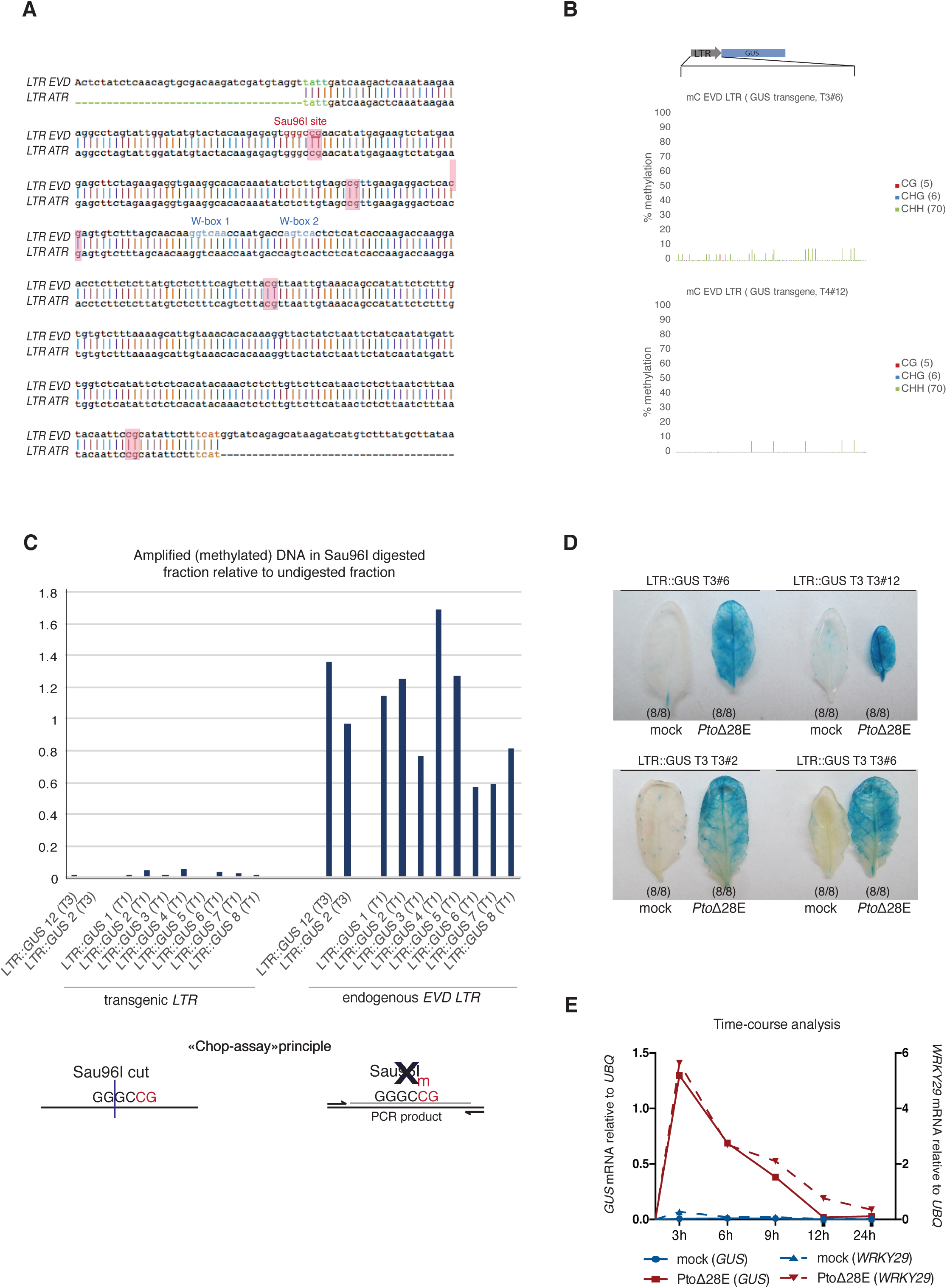
**A.** Alignment between the LTRs of *ATCOPIA93-EVD* and *ATCOPIA93-ATR* showing that they are identical in sequence. The two W-boxes tested in Figure 1 are highlighted in blue; the beginning and end of the LTRs of the COPIA93 family are highlighted in green and orange respectively; the GGGCC sequence in red is the site recognized by Sau96I methylation-sensitive restriction enzyme and underlined is the CG site analyzed by Chop-qPCR in Figure EV1C and Figure 3D. The five CG sites of the LTR are highlighted with pink boxes. **B.** Cytosine methylation analyzed by bisulfite-sequencing at the LTR::*GUS* transgene. Genomic DNA of a pool of LTR::*GUS* transgenic plants (one T3 line -T3#6, upper panel- and one T4 line-T4#12, bottom panel-, four plants, two rosette leaves each) was treated with sodium bisulfite, amplified with primers specific for the LTR contained in the LTR::*GUS* construct and cloned for sequencing (21 to 11 clones-towards the end of the sequence- were analyzed for T3#6 and 12 clones were analyzed for T4#12) as in Fig 1A. The percentage of methylated cytosines is indicated by vertical bars. **C.** Methylation status of the DNA (CG site) at the LTR in LTR::*GUS* transgenic plants by Sau96I Chop-qPCR. DNA from a pool of leaves of single primary transformants and pools of T3 homozygous plants was digested with the methylation sensitive restriction endonuclease Sau96I which recognizes GGNCC sites- here GGGCCG- and is sensitive to methylation of the second C. Digested DNA was quantified by using qPCR with primers spanning a Sau96I restriction site in the LTR. On the left, primers were specific for the transgenic LTR (one primer in the vector): lack of amplification shows unmethylation of the CG site in the transgenic LTR. On the right, as a control, the same DNA was analyzed with primers specific for the endogenous *EVD* LTR (one primer upstream of *EVD*): amplification shows methylation of the CG site in the endogenous 5’ LTR sequence. The signal was normalized to an undigested control. The assay principle is schematized under the graph. **D.** Analysis of additional, independent LTR::*GUS* transgenic lines: representative pictures of leaves infiltrated with water (mock), *Pto* DC3000 deleted of 28 effectors (*Pto*Δ28E) at 2.10^8^ colony-forming unit per ml (cfu/ml) with GUS substrate 24 hours post-infiltration (24hpi). The number of leaves showing this representative phenotype is indicated into brackets. **E.** Additional independent experiment for the time-course analysis of *GUS* mRNA (plain lines) and PTI-marker *WRKY29* mRNA (dashed lines) by RT-qPCR. The experiment was performed as in Fig 1C.

### *AtCOPIA93* reactivation is negatively controlled by DNA methylation during PAMP-triggered immunity

We next analyzed, over the same timeframe, the reactivation of the almost identical endogenous copies of *ATCOPIA93*: *EVD* and *ATR*. The induction of their expression upon bacterial challenge was generally weak in wild-type leaves (Fig 2A and 2B, Fig EV2). By contrast, we observed a consistent and transient induction of *EVD*/*ATR* at 3, 6 and 9 hpi in a *ddm1* hypomethylated background (Fig 2A and 2B left panel, Fig EV2), specifically in response to the *Pto*Δ28E strain. *EVD*/*ATR* transcript levels were also significantly enhanced at 6 hpi in a bacteria-challenged *met1* mutant (Fig 2B, right panel), which is impaired in CG methylation. Together, these data indicate a tight negative control of *ATCOPIA93* induction which is exerted by DNA methylation and is particularly relevant during bacterial challenge when the LTR is activated. Notably, at this developmental stage, mutations in the components of the DNA methylation pathways were not sufficient to enhance *ATCOPIA93* expression in the absence of bacterial stress, in accordance with the transcriptional behavior of the unmethylated LTR::*GUS* fusion which displays weak promoter activity in water-treated plants (Fig 1).

**Figure 2.**
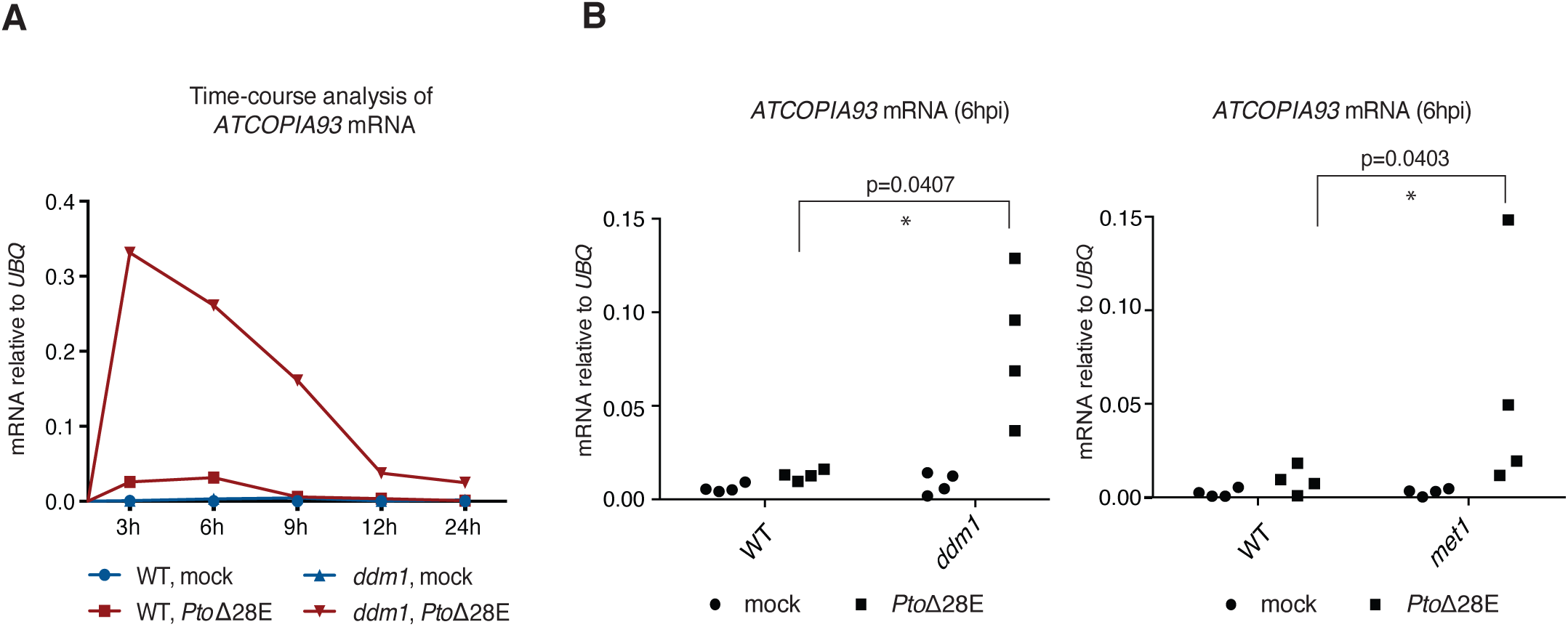
ATCOPIA93 reactivation is negatively controlled by DNA methylation. **A.** Time course analysis of *ATCOPIA93* mRNA by RT-qPCR. Leaves were infiltrated with water (mock), or *Pto*Δ28E bacteria at 2.10^8^ cfu/ml; two similar leaves of three to four plants were pooled by condition and by time-point after infiltration before extracting total RNAs. Values were determined by RT-qPCR and are relative to the expression of the *UBIQUITIN* gene (*At2g36060*). This experiment was repeated twice with similar results and another independent experiment is shown in Fig EV2. **B.** *ATCOPIA93* mRNA analysis in two methylation-defective mutants, *ddm1* and *met1*, at 6 hours post-treatment with either water or *Pto*Δ28E at 2.10^8^ cfu/ml. Material and data were generated as in A. The data points for four independent experiments are plotted. Two-tailed P values were calculated by paired T-test to take into account inter-experiment variability. The variability observed between biological replicates is inherent to the developmental stage (adult leaves) analyzed which often shows differences in the timing and extent of PTI from one experiment to the other (see Fig1C/FigEV1E et 2A/EV2).

**Figure EV2.**
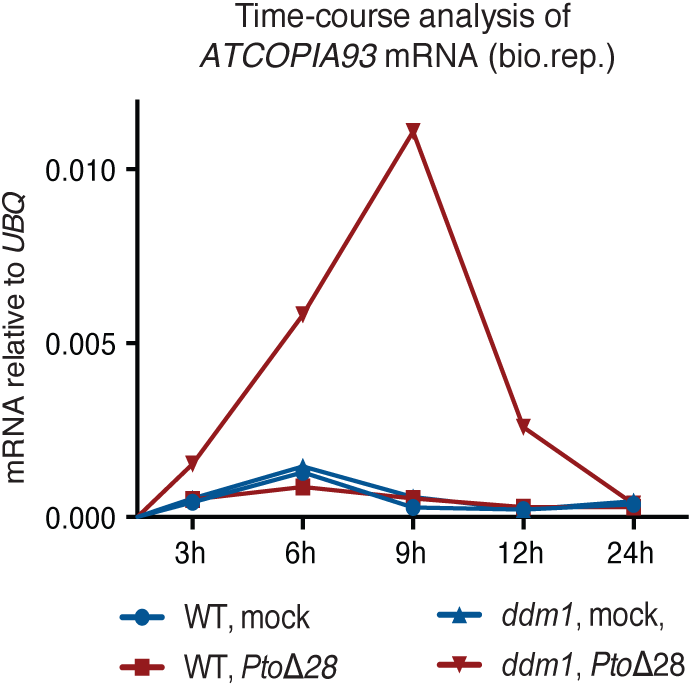
Biological replicate (independent experiment) for the time course analysis of *ATCOPIA93* mRNA by RT-qPCR. The experiment was performed as in Fig 2A.

### *EVD* is marked by H3K27m3 chromatin modification in addition to DNA methylation

Given previous observations in plants and mammals that some loci gain H3K27m3 marks upon their loss of DNA methylation (Mathieu *et al*, 2005; Deleris *et al*, 2012; Weinhofer *et al*, 2010; Reddington *et al*, 2013; Saksouk *et al*, 2014; Basenko *et al*, 2015), presumably mediating “back-up” transcriptional silencing of hypomethylated sequences, we thought that there could be an increase in H3K27m3 marks at *ATCOPIA93*-LTR in *ddm1* plants. To test this possibility, we inspected publically available ChIP-chip and ChIP-seq datasets and found that *ATCOPIA93* LTR is marked by H3K27m3, not only in an hypomethylated mutant *met1* (Fig3A), but also unexpectedly in wild-type plants (Fig 3A, Fig EV3A). However, one limit of ChIP-chip and ChiP-seq datasets is that they do not allow precise determination of the genomic localization of immunoprecipitated repeated sequences, either because of cross-hybridization (ChIP-chip) or the impossibility to accurately map multiple repeated reads (ChIP-seq). To circumvent this problem, we designed specific qPCR primers to discriminate *EVD-LTR* from *ATR-LTR* sequences after ChIP by using upstream genomic sequences (in this experiment, we also included the housekeeping *UBIQUITIN* gene as well as the heterochromatic, H3K9m2-marked transposon *TA3* (Johnson *et al*, 2002) as negative controls, and an intronic and highly H3K27 trimethylated region of *FLOWERING LOCUS C* as a positive control). In addition, we took advantage of the rare SNPs between *EVD* and *ATR* and used pyrosequencing to analyze the immunoprecipitated fragments from the *ATCOPIA93* coding sequence (CDS), where specific primer design is impossible. Interestingly, with both these approaches, we found that there was a strong bias towards *EVD* molecules in the H3K27m3-IPs (Fig 3B, Fig EV3B and Fig 3C, Fig EV3C). This could be due to a positional effect, as the *EVD* sequence is embedded in a larger domain of H3K27m3 that comprises seven adjacent genes (Fig 3A) (*AT5G17080* to *AT5G17140*, coding for either cysteine-proteinases or cystatin-domain proteins —one of them, *AT5G17120,* was included in the ChIP analysis, Fig 3B). Nevertheless, there was less H3K27m3 in the CDS than in the LTR region (Fig 3B, Fig EV3A and EV3B); this likely reflects the previously observed antagonism of DNA methylation/H3K9m2 and H3K27m3 (Mathieu *et al*, 2005; Deleris *et al*, 2012; Weinhofer *et al*, 2010) since DNA/H3K9 methylation levels are higher in the coding sequence of *EVD* than in the LTR (Fig EV3D) (Marí-Ordóñez *et al*, 2013). Finally, we found that H3K27m3 levels were strongly reduced in *clf* plants mutated for the Polycomb-Repressive Complex 2 (PRC2) H3K27 methyltransferase CURLY LEAF (Förderer *et al*, 2016)(Fig 3B and EV3B, EV3A). Interestingly, we observed that H3K27m3 was absent at the transgenic LTR sequence when performing ChIP analysis on LTR::*GUS* transgenic plants (Fig EV3E). This shows that the transgenic LTR is not subjected to deposition of H3K27m3 upon Agrobacterium-mediated transformation. Therefore, the LTR sequence is unlikely to contain Polycomb Response Elements (PREs) i.e. *cis*-localized DNA sequence motifs that would recruit PRC2 as observed for many developmental genes (Xiao *et al*, 2017).

Next, we assessed whether H3K27m3 and DNA methylation could co-exist on the same molecules or whether the detection of both marks in wild type rosette leaves was only reflecting the contribution of different cell types, some marked by H3K27m3 at *EVD* and some by DNA methylation. To distinguish between these two possibilities, we analyzed the DNA methylation status of one representative CG site at the LTR region of *EVD* using a methylation sensitive enzyme assay on H3K27m3-IPed DNA, followed by qPCR (Fig EV1C, “Chop-assay” principle scheme). We observed amplification of the enzyme-treated DNA, comparable to the total input genomic DNA (Fig 3D). Based on this result we can conclude that the *EVD* DNA associated with H3K27m3 is methylated and that DNA methylation does not inhibit H3K27m3 deposition in this region. This true co-occurrence of the two marks, while unexpected and never reported in plant vegetative tissues, was recently observed in the endosperm at pericentromeric transposable elements (Moreno-Romero *et al*, 2016) as well as in mammals where lower densities of CG methylation were found to allow H3K27m3 deposition (Statham *et al*, 2012; Brinkman *et al*, 2012). Accordingly, *EVD* LTR has only five methylated CGs (Fig 1A, Marí-Ordóñez *et al*, 2013), which is low relative to the size of the LTR (roughly 400 bp). Thus, in Arabidopsis vegetative tissues, the co-existence of CG methylation and H3K27m3 can occur, possibly constrained by CG density like in mammals.

**Figure 3.**
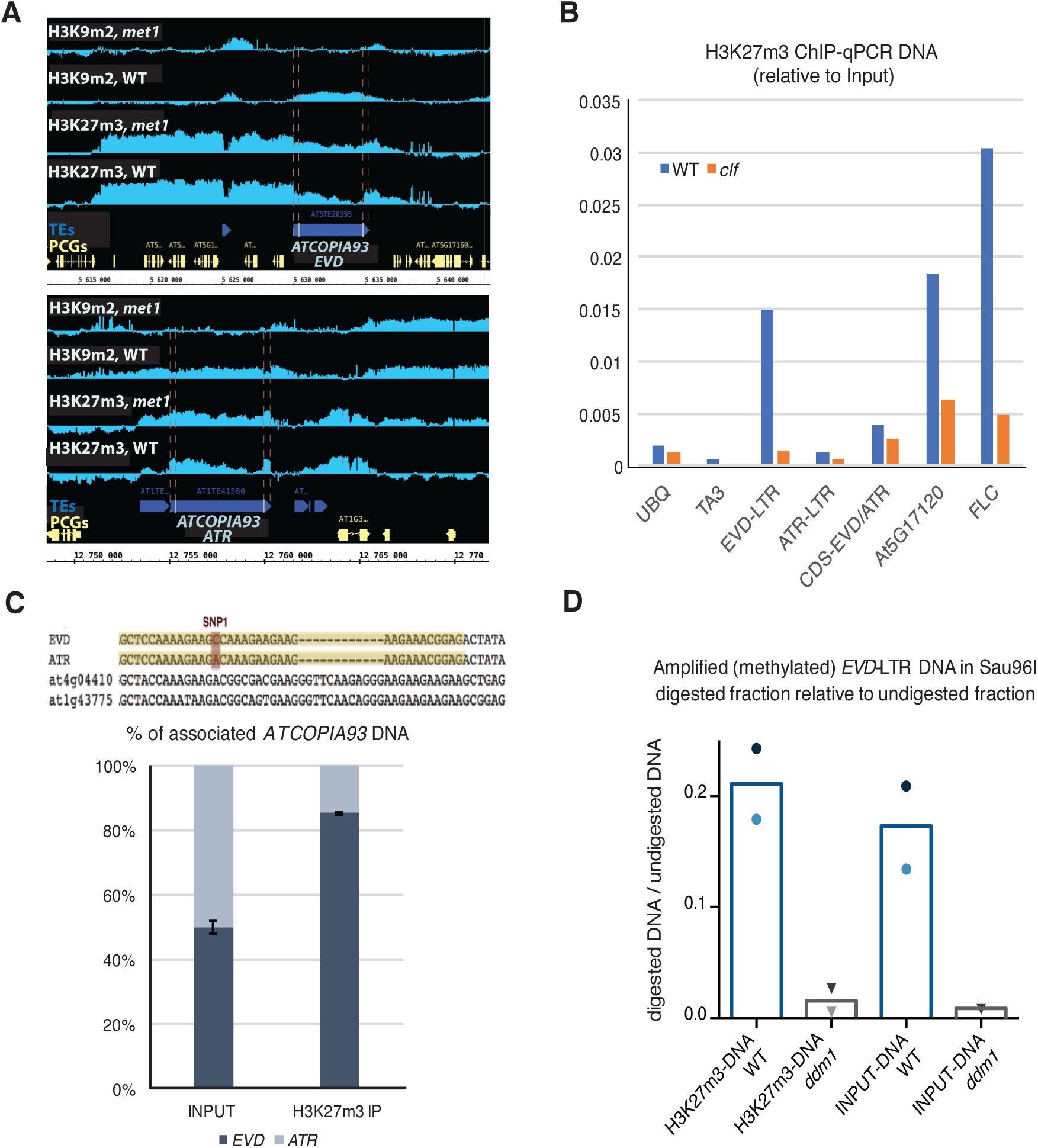
H3K27m3 and DNA methylation coexist at *ATCOPIA93*. **A.** IGB (Integrative Genome Browser) views showing H3K9m2 levels and H3K27m3 levels in WT and *met1* rosette leaves, at *ATCOPIA93 EVD* and *ATR* (ChIP-chip public data, Deleris *et al.,* 2012). Yellow horizontal bars: protein-coding genes; horizontal blue bars: transposable elements. The LTRs are delineated by pink bars. Vertical light blue bars: H3K9m2 signal relative to H3 (two top lanes) and H3K27m3 signal relative to H3 for each probe. **B.** Analysis of H3K27m3 marks at *ATCOPIA93 EVD* and *ATR* by ChIP on rosette leaves, followed by qPCR, in wild-type plants and in *clf* plants mutated for the H3K27 methyltransferase CURLY LEAF. Data were normalized to the input DNA. *ATCOPIA93 CDS* is a region in *ATCOPIA93 GAG* common to *EVD* and *ATR*. *At5g17120* is a region in the protein-coding gene located upstream of *EVD*. *FLC* is a region located in the first intron of *FLOWERING LOCUS C* which shows high levels of H3K27m3 in vegetative tissues and serves as a positive control. *TA3* is a transposon and serves as a negative control. Because of technical variability in the ChIP efficiency, one ChIP experiment is presented here and two other independent experiments are presented in Figure EV3B. ChIPs were performed on a pool of rosette leaves from eight to ten plants/genotype. **C.** Genomic distribution of H3K27m3 marks between *EVD* and *ATR* loci by ChIP-PCR pyrosequencing. Upper panel: depiction of the pyrosequenced region (in yellow) within the *GAG* biotinylated qPCR amplicon obtained after H3K27m3 ChIP-qPCR and purification with streptavidin beads. The position interrogated corresponds to the discriminating SNP between *EVD* (C/G) and *ATR* (A/T). The % indicated represents the % of G (*EVD*, in black bar) or T (*ATR*, grey bar) at that position. The sequencing primer was designed so that other *ATCOPIA93*-derived sequences (divergent and presumably nonfunctional) such as *At4G04410* and *AT1G43775* cannot be amplified and so that the allelic ratio between the two active *ATCOPIA93* copies *EVD* and *ATR* only can be evaluated. To verify this, the qPCR GAG product is also amplified from the Input gDNA as a control where a 50%-50% ratio is expected. For clarity, an average of two experiments performed on two independent Input and ChIPs samples is shown and individual datasets presented in Figure EV3C. **D.** Methylation status of the DNA captured with H3K27m3 by Sau96I Chop-qPCR. H3K27m3 ChIP-DNA from two independent ChIPs was digested with the methylation-sensitive restriction endonuclease Sau96I which is sensitive to the methylation of the second C at the GGGCCG site in the LTR (as in Fig EV1). The values plotted correspond to the ratio between the amount of amplified DNA in the Sau96I digestion and the amount of amplified DNA in the undigested control, as calculated by the formula 2^-^(Ct.digestedDNA-Ct.undigestedDNA) and using primers specific for a region of *EVD* LTR spanning this Sau96I restriction site. Dark and light symbols are used for the first and second experiments respectively. Results show that the WT ChIP-DNA had significantly less digestion compared with the *ddm1* control, thus there was more methylation.

**Figure EV3.**
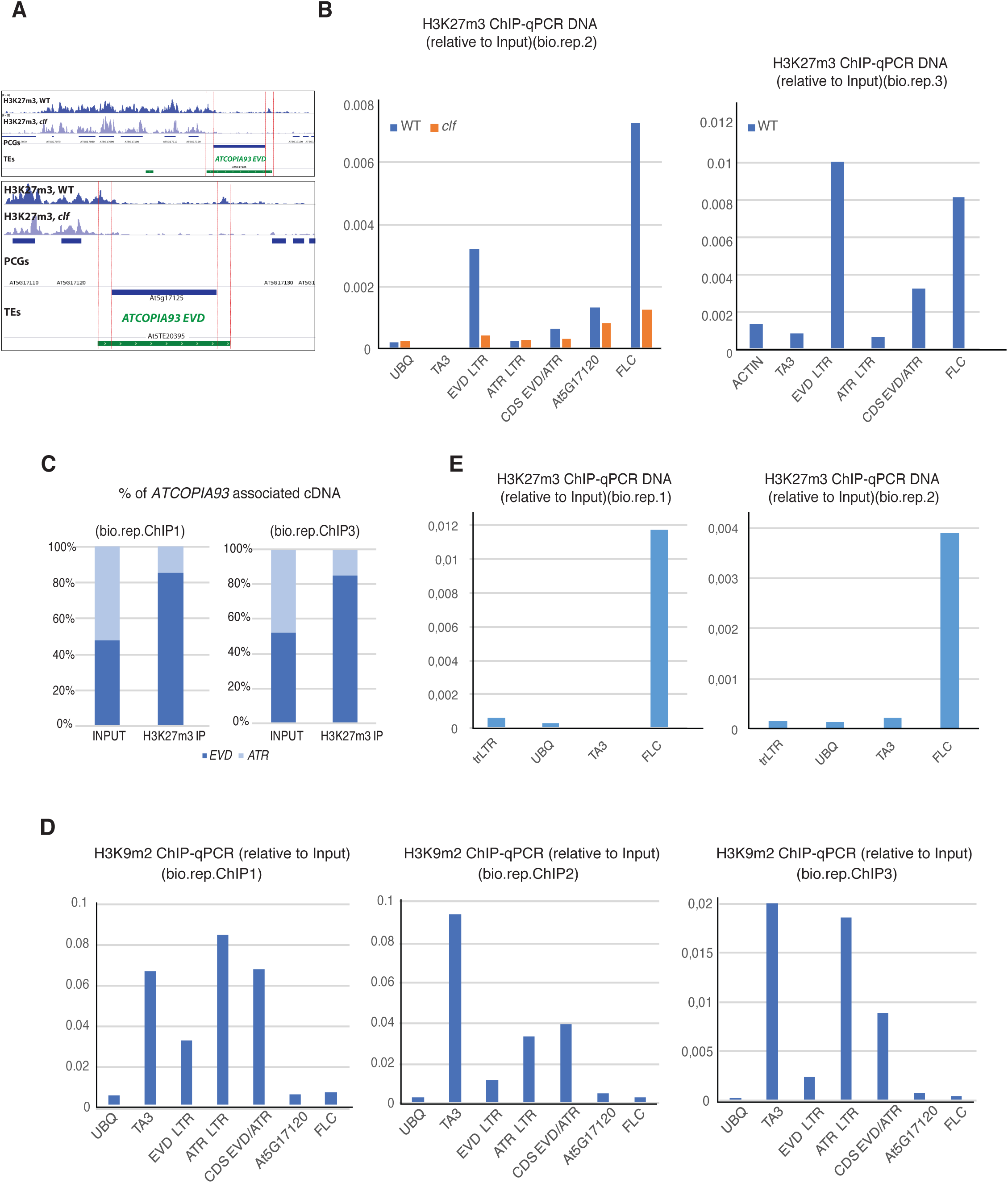
**A.** IGV (Integrative Genome Viewer) showing H3K27m3 levels at *ATCOPIA93 EVD* in wild-type Col and *clf* mutant. Publically available ChIP-seq data (Wang *et al*, 2016) were processed by mapping all reads of the H3K27m3-ChIP libraries including non-unique multi-mappers such as the ones coming from *ATCOPIA93* LTR and CDS. The bottom panel is a close-up view of the top panel. **B.** Additional biological replicates for H3K27m3 analysis at *ATCOPIA93* by ChIP-qPCR. The ChIPs were performed with different amounts of starting material for each batch (bio.rep.) which contributes to explain the differences in ChIP efficiency. Left panel: biological replicate for loss of H3K27m3 marks in *clf*; right panel: additional biological replicate that was used for pyrosequencing of H3K27m3-immunoprecipitated DNA in wild-type plants. **C.** Detail of pyrosequencing replicates on H3K27m3 ChIP-DNA at *ATCOPIA93 CDS*. **D.** Analysis of H3K9m2 marks at *ATCOPIA93 EVD* and *ATR* by ChIP in rosette leaves, followed by qPCR. Data were normalized to the Input DNA. Loci tested are as in Fig 3B. Because of variability in the ChIP efficiency (different amounts of starting material), the three biological replicates are shown separately. **E.** Absence of H3K27m3 marks at the transgenic LTR sequence (“trLTR”) in LTR::*GUS* plants (rosette leaves). qPCRs were performed using primers specific for the transgenic LTR; *UBQ* (*UBIQUITIN*) and *TA3* transposon sequences are used as a negative control for the ChIP while *FLC* (*FLOWERING LOCUS C*) is used as a positive control. Because of variability in the ChIP efficiency (different amounts of starting material), the two biological replicates are shown separately.

### Polycomb-group proteins and DNA methylation exert a dual negative control on *ATCOPIA93* induction during PAMP-triggered immunity

The co-existence of DNA methylation and H3K27m3 at *EVD*-LTR suggests that there is dual control by both PcG- and DNA methylation-mediated silencing on the same molecule, in the same cell type. To test for the functional relevance of PcG silencing at *ATCOPIA93*, we challenged wild-type and *clf* mutant plants with *Pto*Δ28E and monitored *ATCOPIA93* transcript levels by RT-qPCR analyses, using *ddm1* mutant plants as a positive control. Results from these analyses revealed a modest increase of *ATCOPIA93* expression in bacteria-elicited *clf* plants compared to the wild type, though weaker and less reproducible than in elicited *ddm1* plants (Fig 4A, EV4A). Furthermore, by pyrosequencing *ATCOPIA93* cDNA in bacteria-elicited plants, we observed that while both *EVD* and *ATR* are induced in *ddm1*, it was mostly *EVD* that became reactivated in *clf*-elicited mutant background (Fig 4B, Fig EV4B). This is consistent with *EVD* exhibiting comparatively stronger H3K27m3 enrichment than *ATR* in wild-type (Fig 3). The pyrosequencing result also presumably explains, at least partly, the weaker *ATCOPIA93* induction observed in *clf* compared to *ddm1* after bacterial challenge (Fig 4A, EV4A), as the mRNA is almost only contributed by *EVD* in *clf* (Fig 4B). In addition, as anticipated, we observed that in *clf* mutants, where H3K27m3 is reduced, H3K9m2 marks were retained to levels comparable to WT at *EVD* (Fig EV4C, left panel); this presumably contributes to explain lower accumulation of *EVD* transcripts in *clf* than in *ddm1*. In *ddm1* mutants, where H3K9m2 is reduced, H3K27m3 marks were also retained to levels comparable to WT at *EVD* (Fig EV4C, right panel) suggesting that H3K9m2/DNA methylation may exert a stronger repressive effect on *EVD* than Polycomb group proteins.

**Figure 4.**
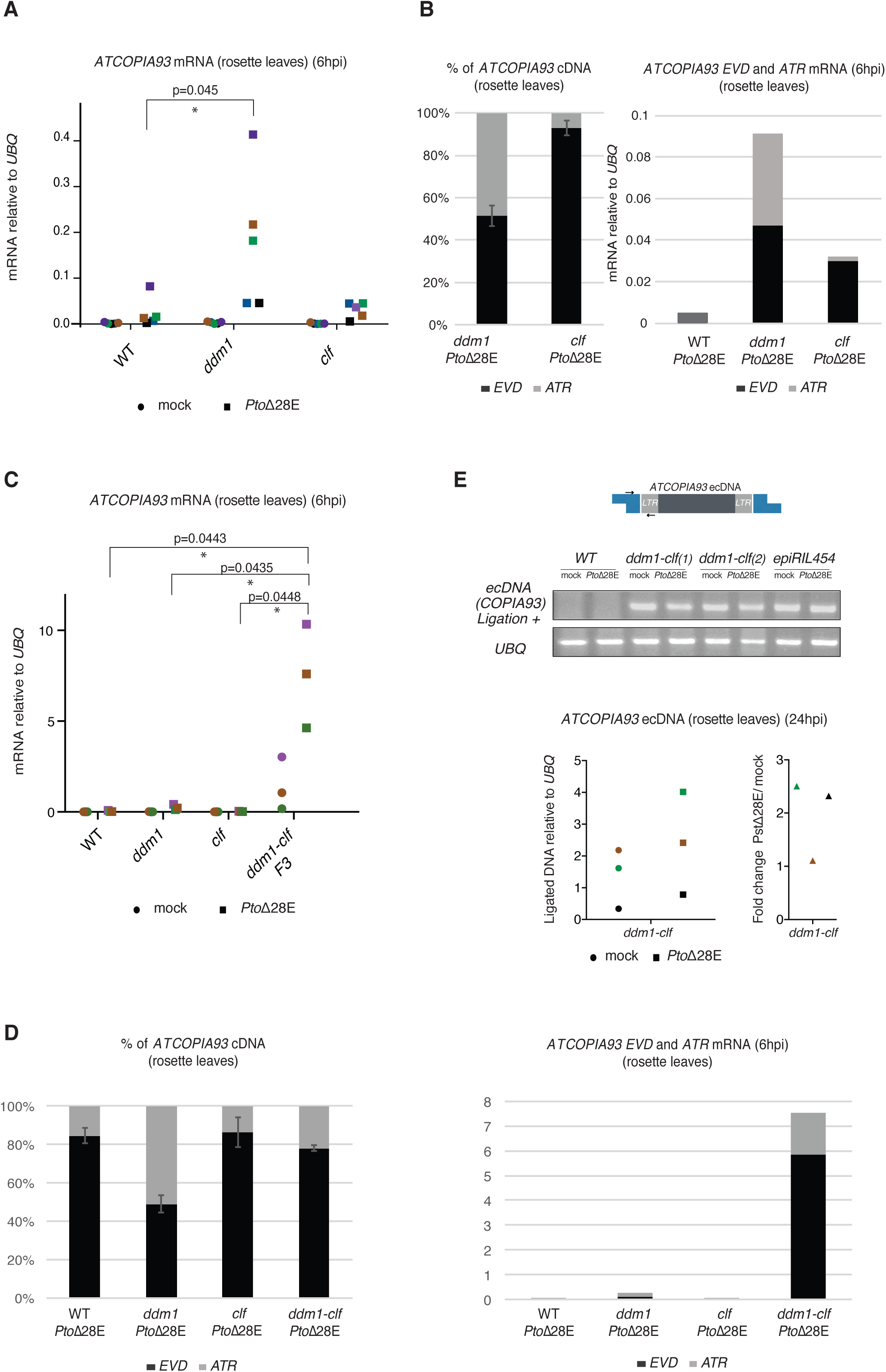
PcG-mediated silencing and DNA methylation exert a dual and differential negative control on the induction of *EVD* and *ATR* during PAMP-triggered immunity. **A.** *ATCOPIA93 mRNA* levels in pools of rosette leaves of DNA methylation mutant *ddm1* and PRC2 mutant *clf* (three to four plants per condition), 6 hours post-infiltration with either water or *Pto*Δ28E bacteria at 2.10^8^ cfu/ml. Values were determined by RT-qPCR and are relative to the expression of the *UBIQUITIN* (*At2g36060*) gene. Five independent experiments were performed and the five corresponding biological replicates are shown and represented by black, blue, green, brown and purple symbols respectively. Two-tailed P values were calculated by paired T-test to take into account inter-experiment variability. The variability observed between biological replicates is inherent to the developmental stage (adult leaves) analyzed which often shows differences from one experiment to the other, in the timing and extent of PTI (see Fig1C/FigEV1E et 2A/EV2). **B.** Left: Qualitative analysis by pyrosequencing of the RT-qPCR products quantified in 4A. The pyrosequenced region and SNP interrogated are the same as in Fig 3. For clarity, the average of three experiments on three of the biological replicates (4A) is shown; the independent replicates are shown individually in Figure EV4B with the corresponding color code. Right: Determination of *EVD* and *ATR* transcripts levels 6hpi with *Pto*Δ28E by integrating *ATCOPIA93* total transcript levels (4A) with pyrosequencing data. Calculations were made by applying the average respective ratios of *EVD* and *ATR* (left panel) to the average RNA values (relative to *UBIQUITIN* and shown in 4A) of the three pyrosequenced biological replicates. Pyrosequencing could not be performed in wild type because of too low amount of *ATCOPIA93* transcript thus the EVD/ATR ratio could not be determined and an intermediate grey color is used. **C.** *ATCOPIA93 mRNA* levels in pools of rosette leaves of *ddm1-clf* double mutants (three to four plants per condition), 6 hours post-infiltration with either water or *Pto*Δ28E bacteria at 2.10^8^ cfu/ml. Values were determined as in A. Three independent experiments were performed and the corresponding biological replicates are shown and represented by purple, green and brown symbols respectively. Due to high values obtained in the double mutants, the scale is different from the scale in A. Two-tailed P values were calculated by paired T-test. **D.** Left: Qualitative analysis by pyrosequencing of the RT-qPCR products quantified in 4C and as in 4B. The average of the values obtained from the three experiments above (4C) is shown; the independent replicates are shown individually in Figure EV4E with the corresponding color code. Right: Determination of *EVD* and *ATR* transcripts levels 6hpi with *Pto*Δ28E by integrating *ATCOPIA93* total transcript levels (4C) with pyrosequencing data. Calculations were made as in 4B. **E.** Top: *ATCOPIA93* linear extrachromosomal DNA (ecDNA) was detected by adaptor-ligation PCR as previously described (Takeda *et al*, 2001; Mirouze *et al*, 2009b) (see scheme) on DNA from various genotypes. Pools of rosette leaves (three to four plants per condition), 24 hours post-infiltration with either water or *Pto*Δ28E bacteria at 2.10^8^ cfu/ml were used. *UBIQUITIN* (*UBQ*) was used to control genomic DNA amounts ; epiRIL”454” (Marí-Ordóñez *et al*, 2013) was used as a positive control for *ATCOPIA93* ecDNA accumulation and two F3 *ddm1-clf* lines were tested. A technical replicate (independent ligation experiment) along with negative controls lacking ligation is shown in Fig EV4F. Bottom: qPCR analysis of the ecDNA using the same primers as above in ddm1-clf mutant. Three biological replicates are shown.

**Figure EV4.**
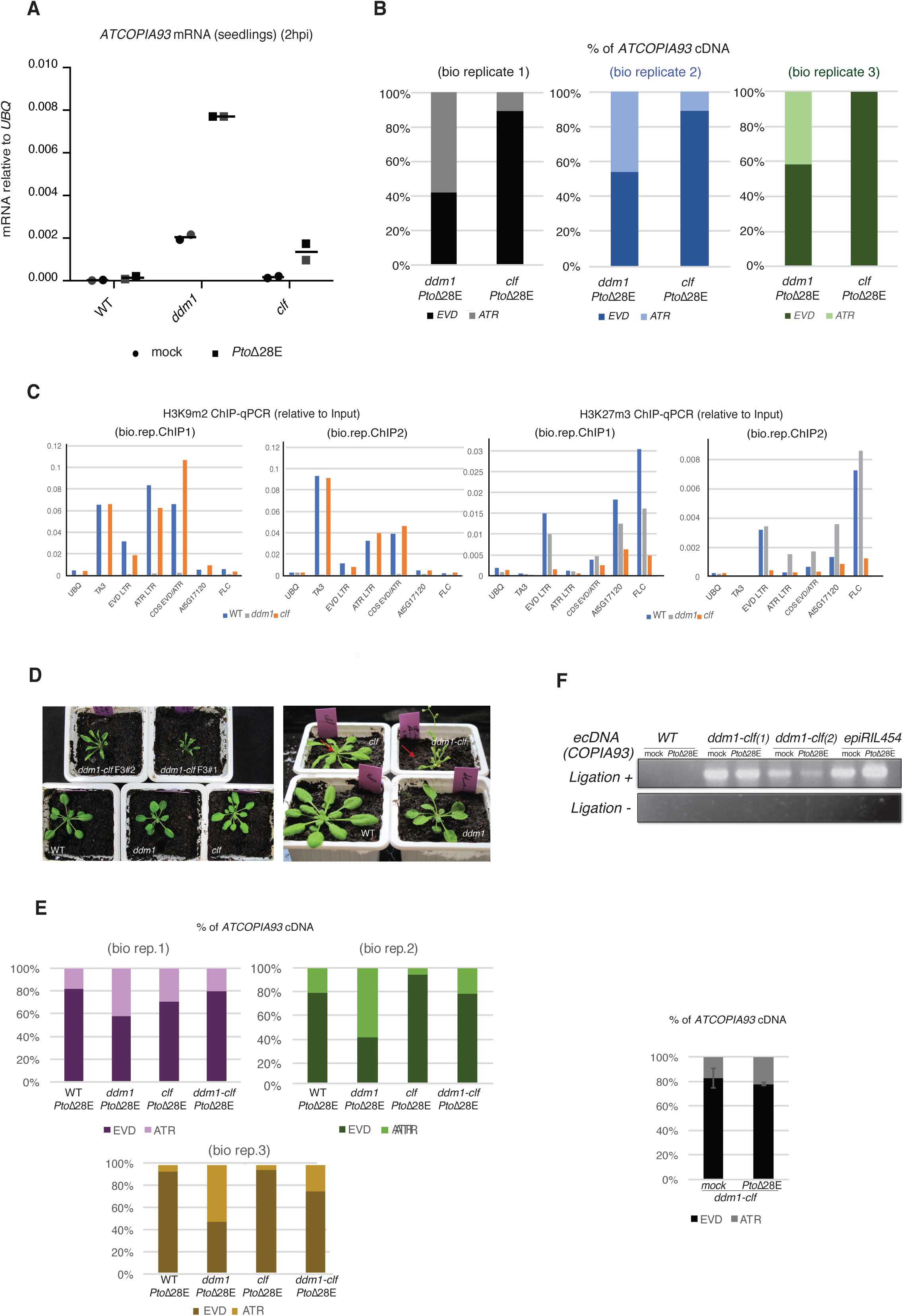
**A.** *ATCOPIA93* mRNA levels in seedlings of *ddm1* and *clf* mutants. About thirty to forty of three week-old seedlings (grown in plates then transferred to liquid medium) were vacuum-infiltrated with either water or a suspension of PtoΔ28E bacteria at 2.10^8^ cfu/ml and collected two hours later (this time-point was determined on the basis of LTR::*GUS* expression in a pilote experiment). Less variability is observed at this stage as shown by the two biological replicates (black and grey symbols). **B.** Detail of the pyrosequencing biological replicates for *ATCOPIA93* cDNA analysis. Each color corresponds to each biological replicate shown in Fig 4A. **C.** Analysis of H3K9m2 and H3K27m3 marks in *clf* and *ddm1* at *ATCOPIA93 EVD* and *ATR* by ChIP in rosette leaves, followed by qPCR. Data were normalized to the Input DNA. Two biological replicates are presented. ChIPs were performed in parallel in WT, *ddm1* and *clf* samples. The analysis of the WT ChIP DNA is already presented in Fig 3B and Fig EV3C (bio.rep.1 and 2) for comparisons between LTR and CDS, and between *EVD* and *ATR*. Here, the *ddm1* and *clf* data were included to show that H3K9m2 marks persist in *clf* mutant (and, as expected, are almost absent in the *ddm1* negative control) and H3K27m3 marks persist in *ddm1* (and are strongly reduced in *clf* as already shown in Fig 3B). **D.** Representative pictures of *ddm1-clf* mutants alongside WT plants and *ddm1* and *clf* single mutants (4.5 weeks-old plants grown in short days). The red arrow indicates that the clf mutants have started to bolt. **E.** Detail of the pyrosequencing biological replicates for *ATCOPIA93* cDNA analysis. Each color corresponds to each biological replicate shown in Fig 4D. In addition, values obtained for mock-treated ddm1-clf double mutants are shown on the right panel. **F.** Technical replicate (independent ligation) for ecDNA detection on the same samples as 4E. The corresponding negative controls lacking ligation (“Ligation-“) were included.

To address the relevance of the double layer of regulation of *EVD* by DNA methylation and H3K27m3, we generated a *ddm1 clf* double mutant (Fig EV4D). In *ddm1 clf* plants, even in the absence of elicitation, *ATCOPIA93* mRNA levels were higher, in average, than in any other genetic background analyzed; in addition, we could observe a significant and quantitatively important increase of *ATCOPIA93* expression upon *Pto*Δ28E challenge (Fig 4C) compared to the wild type and the single mutants. Although it is possible that indirect effects contribute to this striking synergistic effect of *ddm1* and *clf* mutations, these results show that *ATCOPIA93* expression is dually controlled by both DNA methylation and Polycomb-mediated silencing. Pyrosequencing of the cDNA further showed that it was mostly *EVD* that was reactivated in *ddm1 clf* and in *ddm1 clf* -elicited mutant background (Fig 4D, Fig EV4E). This indicates that in the absence of silencing marks *EVD* tends to be more transcribed or expressed than *ATR*, possibly because of position effects, and this balance is shifted towards one TE or the other depending on the present marks in each genetic background.

We next wanted to test whether the high levels of *ATCOPIA93* expression in *ddm1 clf* mutants upon bacterial elicitation could lead to *ATCOPIA93* transposition. Detection of transposition by southern blot and transposon display requires the new insertions to have been clonally inherited during cell division; therefore, these two techniques are not sensitive enough to detect transposition events upon bacterial elicitation of adult leaves, a developmental stage in which cell division has mostly ceased. However, we could clearly detect intermediates of transposition in the form of retrotranscribed, linear, extra-chromosomal (ec) DNA in *ddm1 clf* (Fig 4E, EV4F). We further analyzed the plant DNA at 24 hours post-infiltration with *Pto*Δ28E and could observe an increase in these retrotranscribed forms by quantitative PCR in two biological replicates out of three (Fig 4E, bottom panel). The ecDNA increase observed during the immune response is less pronounced and less consistent than the mRNA increase. This possibly reflects the over-accumulation of the alternative RNA isoform coding for structural TE components (GAG) *versus* the full-length and retrotranscribed *EVD* RNA (Oberlin *et al*, 2017) – excess which is then amplified by the activation of the LTR during immune response. Thus, *ATCOPIA93* potential for transposition is greatly enhanced in the *ddm1 clf* background and further increases during innate immune response.

### *Cis*-regulation of the *RPP4* disease resistance gene by a *ATCOPIA93*-derived, unmethylated soloLTR

The corollary of our findings on *ATCOPIA93* regulation is that the presence of a *ATCOPIA93* LTR in the genome, if deprived of DNA methylation and H3K27m3, can potentially lead to the transcription of downstream sequences, thus potentially affecting the transcription of adjacent genes. We found through a BLAST search, three new *ATCOPIA93*-derived sequences in the genome on the chromosome 3, in addition to the ones that were previously annotated on chromosomes 1, 3 and 4. Interestingly, apart from two copies on chromosomes 1 and 4 (*AT4G04410* and *AT1G43775*, (Mirouze *et al*, 2009)), all other sequences are present in the form of a soloLTR, which is the product of unequal recombination between the LTRs at the ends of a single retroelement (Fig EV5A). The functional W-box 1 was conserved in all of them making them potentially regulatory units responsive to PAMP-triggered immunity (Fig EV5A). Interestingly, transcription was detected in response to various bacterial challenges downstream of soloLTR-1, soloLTR-2 and soloLTR-5 (Fig EV5A). While soloLTR-1 and -2 are located upstream of a pseudogene and an intergenic region respectively, both of unknown function, the soloLTR-5 is embedded in the predicted promoter of the *RPP4* gene, less than 500 bp upstream of the annotated transcriptional start site. *RPP4* is a canonical and functional disease resistance gene that belongs to the *RPP5* cluster on chromosome 4, which is composed of 7 other Toll Interleukin-1 Receptor (TIR) domain-Nucleotide binding site (NBS) and Leucine-rich repeat (LRR) domain (TIR-NBS-LRR genes) (Noel, 1999). Among these genes, *RPP4* was previously shown to confer race-specific resistance against the oomycete *Hyaloperonospora arabidopsidis* isolates Emwa1 and Emoy2 (Van Der Biezen *et al*, 2002). In addition, we observed that *RPP4* was induced during PTI, either triggered by *Pto*Δ28E (Fig EV5B) or by various bacterial and oomycete PAMPs tested at 4 hpi (https://bar.utoronto.ca/eplant/AT4G16860, Tissue and Experiment eFP viewers, Biotic Stress Elicitors, Waese *et al*., 2017). Interestingly, the soloLTR-5 is completely DNA unmethylated and not marked by H3K27m3 nor H3K9m2 (Fig 5A, Ordonnez et al., 2013).

**Figure 5.**
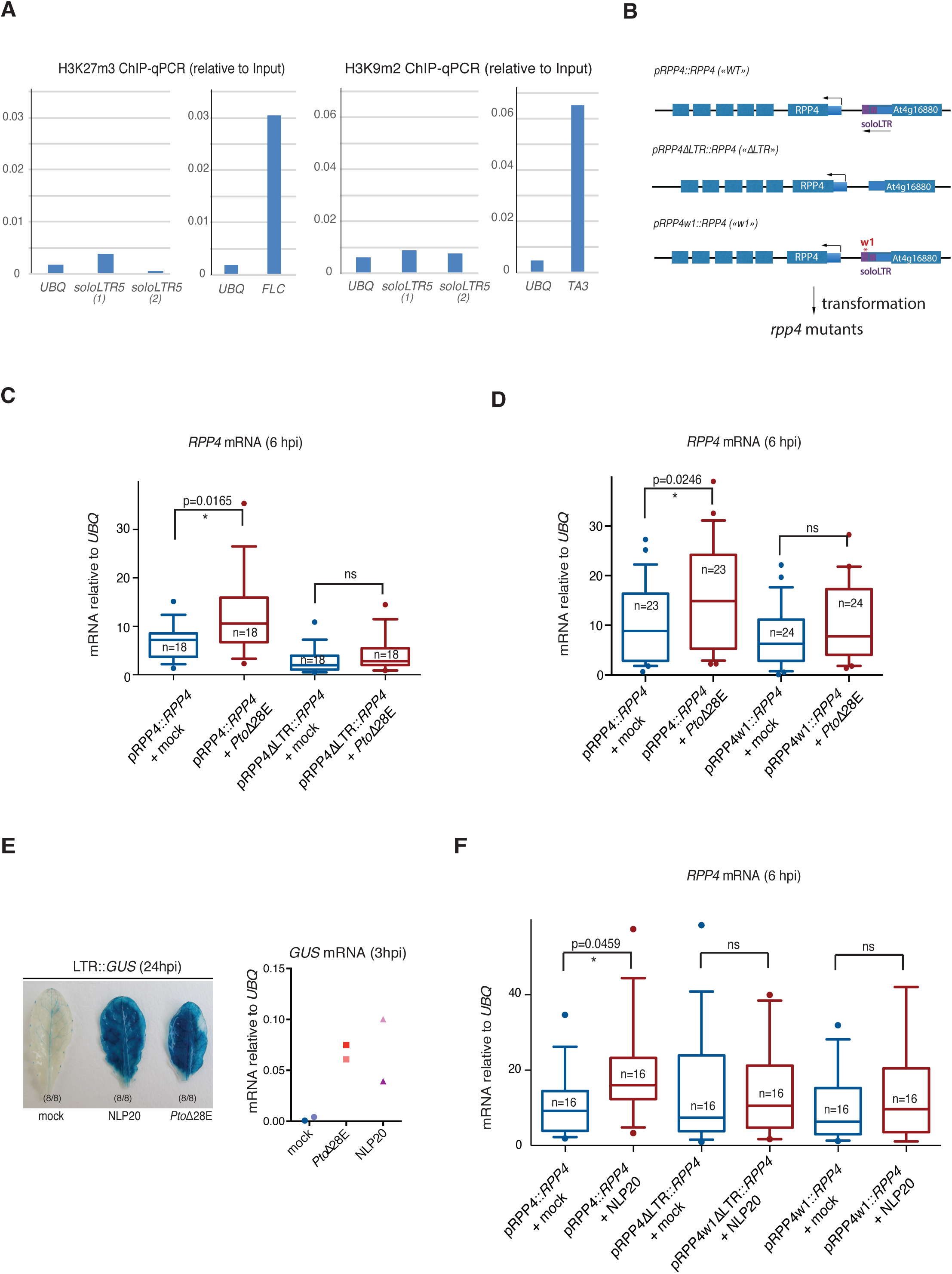
*Cis*-regulation of the *RPP4* immune resistance gene by a *ATCOPIA93*-derived, unmethylated soloLTR. **A.** Absence of H3K27m3 and H3K9m2 marks at the soloLTR-5 (2 primers sets (1) and (2)), upstream of *RPP4*. qPCRs were performed on ChIP-DNA previously analyzed in Fig 3B and Fig EV3C (bio.rep.1) to further validate the epigenetic status inferred from both H3K27m3 and H3K9m2 ChIP-chip (Deleris *et al.,* 2012) and H3K27m3 ChIP-seq data for unique reads (Wang *et al.,* 2016). *UBQ*: negative control for both H3K27m3 and H3K9m2 ChIPs; *FLC and TA3*: positive controls for H3K27m3 and H3K9m2 ChIP respectively. **B.** Depiction of the constructs used to transform the *rrp4* null mutant to assess the impact of LTR mutations on *RPP4* expression. Blue large bars: exons, blue medium bars: transcribed and untranslated regions (UTRs), purple bar: soloLTR-5. **C.** Box plot representing the mRNA levels of *RPP4* in the presence/absence of the soloLTR-5. 18 primary transformants were analyzed for the «p*RPP4*::*RPP4*+ mock» and «p*RPP4*::*RPP4*+ *Pto*Δ28E» datasets and 18 primary transformants were analyzed for «p*RPP4*ΔLTR::*RPP4*+ mock» and «p*RPP4*ΔLTR::*RPP4*+ *Pto*Δ28E» datasets. Mock and *Pto*Δ28E (2.10^8^ cfu/ml) infiltrations were performed on two leaves of each individual transformant that were collected at 6 hours post infiltration (hpi). RNA was extracted for each transformant individually, for each treatment, and analyzed by RT-qPCR to determine the *RPP4* mRNA levels relative to *UBIQUITIN* (*At2g36060*) expression. The values obtained for each primary transformant were plotted. The horizontal line in the box represents the median; the edges of the box represent the 25th and 75th percentiles, the whiskers stretch out to the 10-90 percentile above and below the edges of the box; the symbols (dots) represent the outliers. Two tailed p-values were calculated by unpaired T-test with Welch’s correction. In addition, a non-parametric test (Mann-Whitney) was used and showed the same statistical differences (see Source data for P-value summary). **D.** Box plot representing the mRNA levels of *RPP4* in the presence of the W-box1 or the mutated W-box1 (according to Fig 1E) in the soloLTR-5. Twenty-three primary transformants were analyzed for the «p*RPP4*::*RPP4*+ mock» and «p*RPP4*::*RPP4*+ *Pto*Δ28E» datasets and 24 primary transformants were analyzed for «pRPP4w1::RPP4+ mock» and «pRPP4w1::RPP4+ *Pto*Δ28E» datasets. Analyses were as in C (T-test and non-parametric test showing the same statistical differences, see Source data for P-value summary). **E.** The oomycete PAMP NLP20 induces the same molecular responses as *Pto*Δ28E. Left: Representative pictures of leaves infiltrated with water (mock), 1 μM of NLP20 or effectorless bacteria *Pto*Δ28 at 2.10^8^ cfu/ml as a positive control and incubated with GUS substrate 24hpi (two leaves from three plants per treatment). This result was repeated three times. Right: *GUS* mRNA levels at 3hpi with NLP20 or *Pto*Δ28E. Analyses were performed as in Fig 1. **F.** Box plot representing the mRNA levels of *RPP4* in the presence/absence of the soloLTR-5 and presence/absence of the W-box1, in response to 1 μM of NLP20. Sixteen primary transformants for each construct were analyzed as in C and D (T-test and non-parametric test show the same statistical differences, see Source data for P-value summary).

**Figure EV5.**
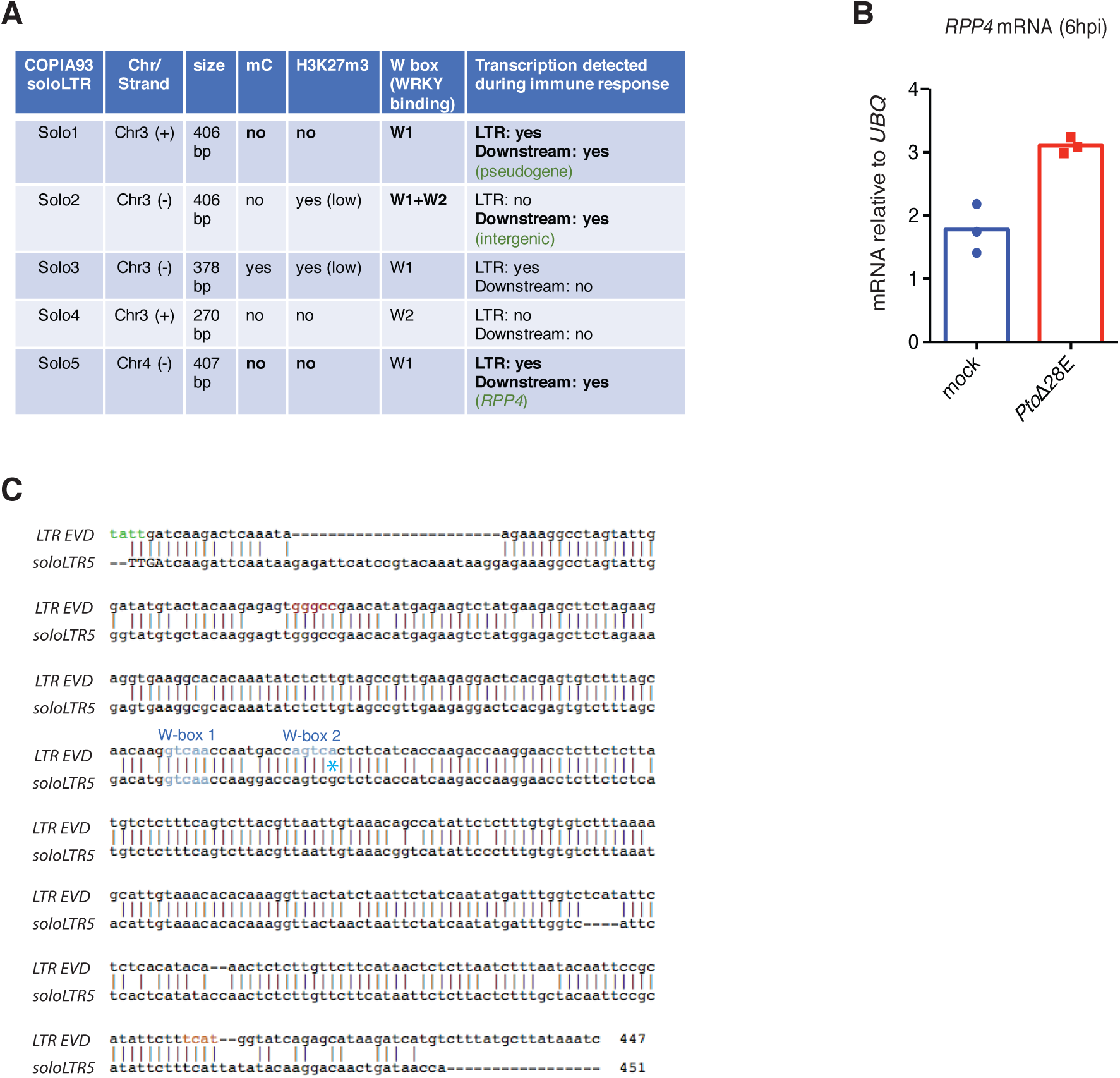
**A.** Summary of the characteristics of the *ATCOPIA93*-derived soloLTRs in Col-0 wild-type plants. chr: chromosome; (+): plus strand; (-): minus strand; mC: methylated cytosines. Methylation status at cytosines and transcription was inferred from inspection of public data at unique reads (http://neomorph.salk.edu/arabidopsis_methylomes/stressed_ath_methylomes.html); trimethylation status of H3K27 was inferred from inspection of both ChIP-chip (Deleris *et al*. 2012) and ChIP-seq data for unique reads (Wang *et al*., 2016). Transcription downstream of the soloLTRs was inspected in plants treated with various bacteria (http://neomorph.salk.edu/arabidopsis_methylomes/stressed_ath_methylomes.html)(Dowen *et al.,* 2012); in addition, *RPP4* induction was observed in *Pto*Δ28E-treated plants (Figure EV5B) and in response to various bacterial and oomycete elicitors (https://bar.utoronto.ca/eplant). **B.** *RPP4* mRNA levels in wild-type plants, at 6hpi with either water or *Pto*Δ28E bacteria. Two similar rosette leaves of three plants were used per condition. Values were determined by RT-qPCR and are relative to the expression of the *UBIQUITIN* gene. Three independent biological replicates are shown. **C.** Alignment between the soloLTR-5 (upstream of *RPP4*) and the *EVD/ATR* soloLTR. The W-box 1 (GGTCAA), which is conserved in both, is depicted in blue.

To test whether the presence of this presumably PAMP-responsive *ATCOPIA93*-soloLTR has a *bona fide* impact on the expression of *RPP4* during PTI and could be co-opted for regulatory functions, we took a loss-of-function approach. We transformed *rrp4* knock-out (KO) mutants with transgenes consisting of the entire *RPP4* genomic region under the control of its native promoter (∼3kb upstream of the TSS) or under the control of the same promoter sequence with a deletion for soloLTR-5 (Fig 5B, “WT” and “ΔLTR” constructs). We further analyzed the primary transformants for *RPP4* expression in response to either water or *Pto*Δ28E treatments and found that the induction of *RPP4* expression upon bacterial challenge was no longer significant in the absence of the soloLTR-5 (Fig 5C). These results provide evidence that the *ATCOPIA93*-derived LTR is required for proper induction of *RPP4* upon bacterial stress. In addition, we wanted to assess whether soloLTR-5 contributes to *RPP4* induction during *Pto*Δ28E elicitation through a W-box motif. We noticed that the W-box 2 present in *EVD/ATR*-LTR is absent in the soloLTR-5 as there is a single nucleotide polymorphism (SNP) in the core motif (GTCA>GTCG) (Fig EV5C). We thus tested the importance of the conserved W-box 1, which we had found earlier to have a partial effect on the induction of the LTR::*GUS* fusion (Fig 1E). To do so, we transformed the *rpp4* KO mutant plants with a construct comprised of the *RPP4* gene under the control of its promoter mutated in the W-box 1 element (Fig 5B, “w1” construct). We found that induction of *RPP4* expression no longer occurred in these “w1” transgenic plants (Fig 5D). These results demonstrate that the LTR responsiveness to bacterial PAMPs contributes to proper induction of *RPP4* and is mediated, at least in part, by a functional W-box element.

Finally, to assess the relevance of this layer of regulation of *RPP4* during immune responses against *H. arabidopsidis* (to which *RPP4* confers race-specific resistance), we treated LTR::*GUS* plants with NLP20, the active peptide of the oomycete PAMP NPP1 that was previously shown to be non-cytotoxic *in planta* (Böhm *et al*, 2014). We found that the LTR::*GUS* fusion was similarly responsive to this oomycete PAMP (Fig 5E), showing that NLP20 induces the same *ATCOPIA93*-LTR regulation as *Pto*Δ28E, in accordance with the fact that the PTI responses induced by unrelated PAMPs largely overlap (Katagiri, 2004; Schwessinger & Zipfel, 2008; Zipfel *et al*, 2006). Importantly, the induction of *RPP4* in response to NLP20 was compromised when the soloLTR was absent or when the Wbox-1 was mutated (Fig 5F), showing that this regulation is relevant during immune response against oomycetes.

Taken together, our results indicate that a *ATCOPIA93*-derived soloLTR has been co-opted during evolution to *cis*-regulate *RPP4* expression during immune responses triggered by unrelated PAMPs.

## Discussion

*ATCOPIA93* has been a widely used model to study plant transposon biology and epigenetics over the last years (Tsukahara *et al*, 2009; Mirouze *et al*, 2009; Tsukahara *et al*, 2012; Marí-Ordóñez *et al*, 2013; Reinders *et al*, 2013; Rigal *et al*, 2016; Oberlin *et al*, 2017). However, with the exception of DNA methylation-defective mutants, the conditions required for the activation of this family, have not been fully explored. Here, we show that the *ATCOPIA93* LTR, in the absence of negative epigenetic control, has the hallmarks of an immune-responsive gene promoter: i) responsiveness to unrelated PAMPs, ii) transient activation upon PAMP elicitation (like early PAMPs-induced genes), iii) suppression of transcriptional activation by bacterial effectors, iv) full dependence on biotic stress-response elements for activation (W-box *cis*-regulatory elements). While, so far, *EVD* transcripts had been only observed in discrete cell types in the absence of DNA methylation (Marí-Ordóñez *et al*, 2013), we show here that in the presence of the adequate signaling and transcription factors, it can be expressed in other tissues, such as adult leaves.

The connection between TE activation and stress response was particularly well-addressed in studies of the *Tnt1* family of transposons in tobacco, which was found to be responsive to various biotic and abiotic stresses (Pouteau *et al*, 1991, 1994; Moreau-Mhiri *et al*, 1996; Mhiri *et al*, 1997; Grandbastien *et al*, 1997). Additionally, different *Tnt* families were induced by distinct biotic challenges and functional analyses further proved that the structural motifs present in the LTR sequences of *Tnt1* families provided specific transcriptional reactivation to specific stresses (Beguiristain *et al*, 2001). Here, we show that one single element can be induced by PAMPs from bacterial and oomycetes pathogens; thus, in the future it will be important to determine whether the two functional W-boxes in *EVD/ATR* LTR are differentially involved in the induction by different pathogens, which would indicate the binding of different transcription factors to the same *ATCOPIA93* LTR. Our findings in Arabidopsis, the primary model plant species for epigenetic analyses, should allow for the investigation of the poorly-understood phenomenon of permissiveness of transposon expression to specific stresses, and test whether this differential permissiveness could be epigenetically regulated, as was previously proposed for *Tnt1* (Grandbastien *et al*, 2005).

The conditional induction of *EVD* is reminiscent of the heat-stress responsive element *ONSEN* (Ito *et al*, 2011; Cavrak *et al*, 2014) and expands the repertoire of Arabidopsis model TEs that can potentially highjack the transcriptional host machinery during stress responses. Thus, *ONSEN* is not a unique case, and many Arabidopsis TEs could exhibit this restricted reactivation pattern, where lack of DNA methylation will result in transposition if compounded by specific stress signaling. This would imply that the “mobilome”—the fraction of TEs with transposition activity— observed in *ddm1* unstressed mutants (Tsukahara *et al*, 2009) is likely to be underestimated. Here, we focused on the somatic regulation of *ATCOPIA93* in leaf tissues—where transpositions would not be mitotically inherited thus are difficult to detect—with the aim to test the impact of this regulation on defense gene regulation. However, our *EVD* mRNA and ecDNA analyses suggest increased transposition potential upon bacterial stress: future studies should address the exciting question of enhanced germinal transposition when wild type and *ddm1* flowers are subjected to PAMP elicitation or infected with pathogens.

One major difference between *ATCOPIA93* and *ONSEN* is that *ATCOPIA93* stress-induced expression in the wild type is more tightly controlled by epigenetic regulation for *ATCOPIA93* than for *ONSEN*. Importantly, we revealed that *EVD*, in addition to be subjected to DNA/H3K9 methylation, is targeted by an additional layer of epigenetic control through PcG silencing, which is generally associated with negative regulation of protein-coding genes and miRNA genes in vegetative tissues (Förderer *et al*, 2016). In spite of the reported antagonist effect of DNA methylation —in particular CG methylation— on H3K27m3 deposition, we found that the two marks could co-occur at the *EVD* 5’LTR in vegetative tissues. To explain this observation, we propose that, like in mammals, a low density of CG methylated sites is permissive to the deposition of H3K27m3, since *EVD* LTR sequence contains only five CGs (i.e. a frequency of 1,2 %)— and also only 6 CHG (i.e. a frequency of 1,4%). Interestingly, other Copia LTRs are also characterized by low CG and CHG frequencies as compared to other genomes segments (Appendix S1A) and we further observed that, in average, the CG and CHG density at Copia LTRs is significantly lower than at Gypsy LTRs (Appendix S1B). The first implication of this low CG and CHG density for the LTRs of Copia elements could be that DNA methylation has a less repressive potential at Copia than at Gypsy elements, supporting the idea that epigenetic control of TE silencing should be studied by taking into account TE class (Underwood *et al*, 2017). On the other hand, this low density of CG methylation could be mechanistically linked to a permissiveness to H3K27m3 deposition (Statham *et al*, 2012; Brinkman *et al*, 2012) and Copia LTRs could thus be more prone to be targeted by H3K27m3 which would compensate for lower repression by DNA methylation. Finally, as for the differential H3K27m3 marking between *EVD* and *ATR,* we propose that it is due to, or at least favored by, a position effect of *EVD* which is embedded within a large H3K27m3 domain. Firstly, this idea is supported by the absence of H3K27m3 at the transgenic LTR. It is possible that another sequence than the LTR is responsible for recruiting PRC2 and for the nucleation of a H3K27m3 domain over the TE; however, one would expect, in this scenario, to observe more H3K27m3 on *ATR* because it shares the same sequence as *EVD* —unless H3K27m3 is strongly antagonized by higher levels of DNA methylation than observed at *EVD.* Secondly, the specific genomic localization of *EVD,* next to PcG targets, as well as the almost complete loss of H3K27m3 at *EVD-LTR* in plants mutated for CLF —which was recently shown to be involved in the spreading phase of H3K27m3 at the *FLC* locus C (Yang *et al*, 2017)— rather support the hypothesis wherein H3K27 trimethylation at *EVD* is due to or at least favored by spreading from the neighboring genes marked by H3K27m3. Future studies should identify and delineate TEs that can potentially recruit PRC2 in *cis* from the ones that become H3K27m3-marked due to an insertional effect, and the ones that exhibit both characteristics.

Importantly, we showed that PcG silencing is functional at *EVD ATCOPIA93.* The modest effect of the sole H3K27m3 loss initially seen on total *ATCOPIA93* expression is presumably due to the specific *EVD* control by PcG in wild-type and the functional redundancy of H3K27m3 with DNA methylation which we then revealed in the double *ddm1 clf* mutant. In the last years, there has been increasing evidence supporting a role of PcG in TE silencing. One of them is the recent discovery of negative regulators of PcG, such as ANTAGONIST OF LIKE HETEROCHROMATIN PROTEIN 1 (ALP1), which is encoded by a domesticated transposase (Liang *et al*, 2015) and which is likely evolved to protect TEs from Polycomb-mediated silencing. In addition, cooperation between DNA methylation and PcG had been previously observed —in the form of functional redundancy— in naturally hypomethylated plant cell types such as the plant endosperm (Weinhofer *et al*, 2010) or chemically-hypomethylated mammalian embryonic stem cells (Walter *et al*, 2016). Our study further demonstrates the relevance of Polycomb-mediated silencing in vegetative tissues at a locus where both DNA methylation and H3K27m3 coexist. Interestingly, this negative control of *EVD* by PcG adds an additional epigenetic layer of restriction specifically at the functional *ATCOPIA93* member (which is also less DNA methylated than its pericentromeric counterpart *ATR*), and presumably limits its somatic transposition while the corresponding soloLTR in the *RPP4* promoter is activated for proper gene regulation during immune response. Notably, this double and differential mC/H3K27m3 marking could allow for unique members of one TE family to be differentially regulated, in particular at discrete stages of development since DNA methylation and PcG are not equivalent in their lability. This might provide the family members with different and discrete windows of opportunities to be expressed and transpose in some restricted tissues, and account for a not-yet appreciated strategy of TEs to adapt to their host.

Interestingly, the *ATCOPIA93-*LTR::*GUS* fusion was consistently found to be unmethylated in various transgenic lines in the wild-type Col-0 background, concordant with LTR::*GUS* reactivation in response to *Pto*Δ28E. This lack of *de novo* methylation upon transformation may be explained by weak LTR transcriptional activity in untreated plants thus preventing expression-dependent RNA-directed DNA Methylation (Fultz & Slotkin, 2017) and/or by low levels of CHH methylation/siRNAs at *ATCOPIA93* (Fig 1A) (Mirouze *et al*, 2009; Marí-Ordóñez *et al*, 2013)), thus preventing identity-based silencing *in trans* (Fultz & Slotkin, 2017). Similarly, in wild type plants, the *ATCOPIA93*-soloLTRs appear usually to be constitutively unmethylated (Fig EV5A), in particular the soloLTR-5 described in detail here. This absence of methylation allowed us to test for a role of the *ATCOPIA93* LTR as a “fully competent” transcriptional module in immunity, *i.e.*, not masked by DNA methylation. This is a different role from the one previously described as an epigenetic module, interfering negatively with downstream expression, when the LTR was artificially methylated in *trans* by siRNAs produced by *EVD* after a burst of transposition in specific epiRIL lines (Marí-Ordóñez *et al*, 2013). The latter results may provide an explanation for the peculiar epigenetic control of *EVD*, which is almost exclusively controlled by CG methylation (Fig1A, Mirouze *et al*, 2009), although it belongs to an evolutionary young family of TEs: the preferred targets of POLYMERASE V, siRNAs and the RNA-directed DNA methylation (RdDM) pathway (Zhong *et al*, 2012). We propose that the low levels of *EVD* LTR siRNAs, which could methylate the soloLTR-5 in *trans* if present in larger quantities, could be the result of evolutionarily selection, so that soloLTR-5 remains unmethylated and proper immune response can be properly activated. This possible selection of an unmethylated soloLTR — combined with the loss of one of the two W-box—, could thus be seen as part of a *RPP4* promoter maturation process towards well-balanced *cis*-regulation since sufficient expression of disease resistance genes is required to mediate immunity but their overexpression results in significant fitness costs (Lai and Eulgem 2017). In this respect, since the soloLTR-1 presents the same characteristics as the soloLTR-5 (absence of methylation, only one W-box, downstream transcription detected during immune response), it would be interesting in the future to functionally characterize the nearby pseudogene and test for its role in immunity.

Previous genome-wide studies have suggested that methylated TEs have a genome-wide repressive effect on nearby gene expression —correlated with their proximity to the gene— (Hollister *et al*, 2011; Wang *et al*, 2013) and also that, upon pathogen stress, transcription of immune-responsive genes could be coupled to the dynamic methylation state of the proximal TEs (Dowen *et al*, 2012). Whether the selection/presence of a constitutively unmethylated TE sequence, *a fortiori* a soloLTR which contains *cis*-regulatory motives, in the promoter of PAMP-responsive genes is a general mechanism that contributes to innate immunity is an exciting question which deserves further exploration. In a first attempt to generalize our findings, we searched for and found dozens of immune-responsive genes whose promoters overlap with a TE fragment containing one LTR or an LTR-derived sequence (Appendix Figure S2A and Appendix Table). Interestingly, most of them (67%) were unmethylated (Appendix Figure S2B) and they contained the genes that were the most induced by PAMPs; in addition, they seemed more induced in average compared to the group with a methylated LTR in their promoter (Appendix Figure S2C and Appendix Table). It would certainly be too simplistic to draw conclusions from these correlations between TE methylation status and downstream gene induction during immune response as the number of genes analyzed here is relatively low (and different between the two groups), and many other parameters have to be taken into account : presence/absence of *cis*-regulatory elements in the TE, strength of the promoter, possible differential impact of different methylation contexts on transcription, localization of DNA methylation within the promoter. Nonetheless, the genes we have identified are potential candidates for further studies to generalize the phenomenon we have uncovered for *RPP4*. These studies will certainly benefit from currently arising techniques such as epigenome editing which should allow in the future to methylate or demethylate specifically a particular/discrete TE to test for its impact on nearby gene transcription.

Transposable elements have been proposed to contribute not only to the diversification of disease resistance genes, which are among the fastest evolving genes, but also, following their diversification, to the evolution of their *cis*-regulation, as part of their maturation process (Lai & Eulgem, 2017). Compelling evidence exists for the latter role (Hayashi & Yoshida, 2009; Tsuchiya & Eulgem, 2013; Deng *et al*, 2017; reviewed in Lai & Eulgem, 2017). In the present study, we have brought another demonstration for the *cis*-regulatory role of TEs, and for the first time we have linked the co-option of a soloLTR for proper expression of a functional disease resistance gene (*RPP4*) and the responsiveness of the corresponding full-length retroelement (*COPIA93 EVD/ATR*), through its LTR, during basal immunity. TEs have been long thought to be a motor of adaptive genetic changes in response to stress (McClintock, 1984). The link we established between responsiveness of a retroelement to biotic stress and its co-option for regulation of immunity provides experimental support to McClintock’s early model where TEs play a role in the genome response to environmental cues. Future studies should address the extent of the TE repertoire with such restricted expression, in the light of recent findings showing that disease resistance gene clusters are among the most frequent transposition targets observed in nature (Quadrana *et al*, 2016).

## Materials & Methods

### Plant material and growth condition

Plants were grown at 22°C with an 8h light/16h dark photoperiod (short days) and experiments were generally performed on 4.5 to 5-week-old rosette leaves. Apart from Figure 3, where plants were analyzed in the absence of treatment, plants were infiltrated with a syringe with either water (“mock”), synthetic flg22 peptide (Genescript) at 1µM concentration or a suspension of bacteria as described below, always at the same time in the morning (between 10 and 11.30am, depending on the number of plants to infiltrate). Plants were then covered with a clear plastic dome until tissue harvest to allow high humidity (Xin *et al*, 2016). For Figure 4B, 3.5-week-old seedlings grown on MS plates were transferred to MS liquid medium for at least 24h then infiltrated with either water or a suspension of bacteria and transferred back to light for 2 hours.

### Mutant lines

We used the *met1-3* allele (Saze *et al*, 2003), the *ddm1-2* allele (Vongs *et al*, 1993) and the *clf-29* allele (Bouveret, 2006). For Figures 1 and 2, first generation homozygous *met1-3* and *ddm1-2* mutants were genotyped and used for analysis; for Figure 4, second generation *ddm1*-2 homozygous mutants were used. Double *ddm1 clf* mutants were generated by crossing the above-mentioned mutants and experiments were performed on F3 progenies.

### Generation of transgenic lines

#### LTR::*GUS* transgenic lines

EVD-LTR was cloned into a pENTR/D-TOPO vector, then recombined in a pBGWFS7 binary destination vector, upstream of the GUS sequence. LTR::*GUS* constructs were transformed in the Col-0 accession by standard Agrobacterium-mediated protocol (Clough & Bent, 1998). Primary transformants were selected with Basta herbicide. Three lines were selected on the basis of 3:1 segregation of the transgene (single insertion) and brought to T3 generation (#2, #6, #12) where the transgene was in a homozygous state and all four lines behave similarly as for GUS expression. Most experiments were performed on stable T3 lines (#12) homozygotes for the LTR::*GUS* transgene, and some in their progeny (T4) after checking that the absence of DNA methylation persisted. The mutations in W-boxes 1 and 2 were generated by chimeric PCR and the resulting mutated LTR sequences introduced in a pENTR vector by overlapping PCR and the mutated LTRs cloned in pBGWFS7. Experiments were performed on individual primary transformants for wild type and mutated constructs. Transgenic plants were sequenced to verify the presence of the mutations at the *LTR* transgene.

#### p*RPP4::RPP4* transgenic lines

For the “WT” construct, a 3kb sequence upstream of RPP4 predicted TSS was cloned in pENTRD-TOPO; for the “ΔLTR” construct, the same 3kb sequence minus the soloLTR-5 was synthesized and cloned in the same vector; for the “w1” construct, site-directed mutagenesis was used on the “WT” pENTRD-TOPO. A fragment corresponding to the *RPP4* gDNA (with introns and UTRs) was then amplified from wild-type plants and cloned after the RPP4 promoter in the three different “WT”, “ΔLTR” and “w1” pENTRD-TOPO vectors using restriction enzymes. The resulting vectors were recombined with pH7WG and introduced in *rpp4* knock-out mutants (SALK_017521). Primary transformants were selected on hygromycin and analyzed individually.

### Bacterial strains and preparation of inocula

The bacterial strains used are *Pseudomonas syringae pv tomato Pto* DC3000 (“*Pto*”) and a non-pathogenic derivative of *Pto* DC3000 in which 28 out of 36 effectors are deleted (Cunnac *et al*, 2011), referred to here as “*Pto*Δ28E”. Bacteria were first grown on standard NYGA solid medium at 28°C with appropriate selection, then overnight on standard NYGB liquid medium. Bacteria were pelleted and washed with water twice. Suspensions of 2.10^8^cfu/ml were used for *Pto*Δ28E except for Figure 1C where suspensions of 1.10^7^cfu/ml were used to compare to a same inoculum of *Pto* DC3000.

### Histochemical GUS staining

GUS staining was performed as in (Yu *et al*, 2013). Briefly, leaves were placed in microplates containing a GUS staining buffer, vacuum infiltrated three times during 15 min, and incubated overnight at 37 °C. Leaves were subsequently washed several times in 70% ethanol.

### SDS-PAGE and Western blotting

Leaf total protein extracts were obtained by using the Tanaka method (Hurkman & Tanaka, 1986), quantified by standard BCA assay and 100 µg were resolved on SDS/PAGE. After electroblotting the proteins on a Polyvinylidene difluoride (PVDF) membrane, GUS protein analysis was performed using an antibody against the GFP since pBGWFS7 contains a GUS-GFP fusion and the anti-GFP antibody (Clontech #632380) was more specific, and stained with a standard coomassie solution to control for equal loading.

### DNA extraction and bisulfite conversion

DNA extraction and bisulfite conversion was performed as in Yu *et al*, 2013 except that the DNA was not sonicated before bisulfite conversion and 15 to 22 clones were analyzed per experiment.

### RNA extraction and qRT-PCR analyses

Total RNA was extracted using RNeasy Plant Mini kit (Qiagen or Macherey-Nagel). One µg of DNA-free RNA was reverse transcribed using qScript cDNA Supermix (Quanta Biosciences) and either oligo(dT) and random hexamers mix or a transcript-specific primer for GUS mRNA analysis. cDNA was then amplified in RT q-PCR reactions using Takyon SYBR Green Supermix (Eurogentec) and transcript-specific primers on a Roche Light Cycler 480 thermocycler. For each biological replicate, two or three technical replicates were averaged when the qPCR corresponding values were within 0.5 cycles. Expression was normalized to *UBIQUITIN (At2g36060)* expression. In addition, for Figure 4A, two reverse-transcription reactions were performed for each biological replicate – in particular, in order to obtain enough cDNA for pyrosequencing- and qPCRs technical replicates averaged. The PCR parameters are: 1 cycle of 10 minutes at 95°C, 45 cycles of 10 s at 95°C, 40 s at 60°C.

### Chromatin immunoprecipitation and ChIP-qPCR analyses

ChIPs were performed as in (Bernatavichute *et al*, 2008), starting with 0.3 g to 1 g (per ChIP) of adult leaves that were previously crosslinked by vacuum-infiltration of a 1% formaldehyde solution. Antibodies against H3K27m3 and H3K9m2 are from MILLIPORE (07-449) and ABCAM (ab1220) respectively. Two µl of a 1:10 dilution of the IP was used for qPCR. The PCR parameters are: 1 cycle of 10 minutes at 95°C, 45 cycles of 10 s at 95°C, 40 s at 60°C.

### Methylation-sensitive enzyme assay (“Chop-assay”)

Two hundred ng of gDNA (Fig EV1) or 10 to 20 ng of ChIP-DNA (10 ng for Input DNA) (Fig 3) was digested overnight at 37C with 1 µL or 0,5 µl respectively of Sau96I enzyme (Thermoscientist FD0194). As Sau96I cannot be heat-inactivated, DNA was then purified with a clean-up column (Macherey Nagel nucleospin column) (Fig 1A) or, when the amount of material was limited (Fig 3) by standard phenol-chloroform extraction using glycogen to precipitate the DNA. DNA was eluted in 20 µl of water or pellets were resuspended in 20 µl of water; qPCR were performed using 0,3ul and primers that amplify an amplicon spanning the Sau96I site. The same amount of the corresponding non-digested DNA was used for qPCR as a control and to normalize the data.

### Pyrosequencing

*ATCOPIA93* DNA (ChIP-DNA, cDNA or gDNA as a control) was amplified with a biotinylated (forward) primer in the same region where RNA levels were analyzed and containing a SNP between *EVD* and *ATR*; the biotinylated PCR product (40 µl reaction) was pulled down with streptavidin beads (sigma GE17-5113-01) and the sense biotinylated strand sequenced with a Pyromark Q24 (Qiagen) on the sequencing mode. Input DNA was used as a control for equal contribution of each SNP. Analysis and quality check of the peaks were done with the Pyromark Q24 companion software which delivers pyrograms indicating the % of each nucleotide at the interrogated SNP. These percentages were directly plotted for each biological replicate in the Extended Views and averaged for clarity of presentation in the main figures.

### Linear ecDNA detection

Detection of linear ecDNA of ATCOPIA93 was performed as previously described (Takeda *et al*, 2001; Mirouze *et al*, 2009b). Briefly, genomic DNA extracted by a standard CTAB protocol and treated with RNAseA. One hundred to 200 ng of gDNA was ligated overnight at 16°C to adaptors (which were generated by annealing P275 and P276 oligos described in the Primers Table S2) and using a T4 DNA ligase. PCRs (nested PCRs for gel analysis using oligos P277-P279 then P278-P279) and qPCRs (P278-P279) were performed on a 20x dilution of the ligation.

### Statistical tests

Statistical tests were performed using GraphPad Prism version 6.04 for Windows.

## Authors contributions

J.Z., A.D., A.Y., J.W. performed the experiments, A.D. and J.Z. analyzed the data; D.A. and J.D. performed bioinformatic analyses; A.D, J.Z. and L.N. designed the experiments; A.D. conceived the study and wrote the manuscript.

## Acknowledgments

We thank the members of the Navarro Lab for their input and discussions as well as the Bourc’his and Felix Labs for their help with pyrosequencing, M. Mirouze for sharing the protocol for *ATCOPIA93* linear ecDNA detection, T. Thorsten for providing the NLP20 peptide, A. Collmer for providing the PtoΔ28E mutant strain, D. Bouyer and L. Quadrana for discussions, H. Keller for valuable comments, M. Greenberg and M. Boccara for critical reading of the manuscript and discussions.

## Bullet points

- The unmethylated *ATCOPIA 93* LTR::*GUS* fusion behaves like an immune-responsive gene
- The corresponding endogenous TE “*EVD*” is dually controlled by DNA methylation and Polycomb group proteins during the immune response
- A derived *ATCOPIA93* soloLTR is naturally unmethylated and is required for proper induced expression of the *RPP4* disease resistance gene during plant defense
- We established a link between the controlled responsiveness of a retrotransposon to biotic stress and the co-option of its LTR for plant immunity.

## BlurbText

Immune-responsive *EVD* LTR-retrotransposon is dually controlled by DNA methylation and Polycomb-group proteins. This restricts *EVD* expression and its transposition potential while the corresponding, unmethylated soloLTR in the *RPP4* promoter is activated for proper gene regulation during plant defense.

